# High-resolution structures illuminate key principles underlying voltage and LRRC26 regulation of Slo1 channels

**DOI:** 10.1101/2023.12.20.572542

**Authors:** Gopal S. Kallure, Kamalendu Pal, Yu Zhou, Christopher J. Lingle, Sandipan Chowdhury

## Abstract

Multi-modal regulation of Slo1 channels by membrane voltage, intracellular calcium, and auxiliary subunits enables its pleiotropic physiological functions. Our understanding of how voltage impacts Slo1 conformational dynamics and the mechanisms by which auxiliary subunits, particularly of the LRRC (Leucine Rich Repeat containing) family of proteins, modulate its voltage gating remain unresolved. Here, we used single particle cryo-electron microscopy to determine structures of human Slo1 mutants which functionally stabilize the closed pore (F315A) or the activated voltage-sensor (R207A). Our structures, obtained under calcium-free conditions, reveal that a key step in voltage-sensing by Slo1 involves a rotameric flip of the voltage-sensing charges (R210 and R213) moving them by ∼6 Å across a hydrophobic gasket. Next we obtained reconstructions of a complex of human Slo1 with the human LRRC26 (γ1) subunit in absence of calcium. Together with extensive biochemical tests, we show that the extracellular domains of γ1 form a ring of interlocked dominos that stabilizes the quaternary assembly of the complex and biases Slo1:γ1 assembly towards high stoichiometric complexes. The transmembrane helix of γ1 is kinked and tightly packed against the Slo1 voltage-sensor. We hypothesize that γ1 subunits exert relatively small effects on early steps in voltage-gating but structurally stabilize non-S4 helices of Slo1 voltage-sensor which energetically facilitate conformational rearrangements that occur late in voltage stimulated transitions.

## Introduction

The Slo1 channel, alternately known as BK or MaxiK, underlies the large conductance, voltage and calcium regulated K^+^ selective currents^1–5^, that play key roles in diverse physiological processes such as neurotransmitter release and muscle contraction^6,7^. In higher organisms, the Slo1 channel associates with members of multiple families of auxiliary subunits^8,9^, which can differentially influence its voltage-dependent gating, calcium sensitivity and pharmacology. For instance, the product of the KCNMB2 gene, termed the β2 subunit, confers fast inactivation characteristics on Slo1 currents^10–12^ and reduces its sensitivity to the scorpion peptide toxin, charybdotoxin^13^. The γ subunits, consisting of members of a LRRC (Leucine Rich Repeat Containing) family of single pass transmembrane proteins, form a structurally distinct family of Slo1 auxiliary subunits. The founding member of this family of regulatory subunits, LRRC26 or γ1, dramatically shifts the conductance-voltage (GV) relationship of Slo1 in the hyperpolarizing direction^14^, in nominally calcium free conditions. As a result, in cells such as secretory epithelia where Slo1 is partnered with γ1, these complexes contribute to K^+^ flux under unstimulated, resting conditions^15–17^. A paralogous subunit, LRRC52 (or γ2), associates with Slo1 channels in cochlear inner hair cells^18^ and also with Slo3 channels in mammalian sperm contributing to KSper currents which are critical for sperm capacitation^19^. Evolutionarily related LRRC55 and LRRC38 (γ3 and γ4 respectively) proteins, however, modulate Slo1 gating more modestly^20^ and it remains unclear whether they are *bona fide* auxiliary subunits of Slo1.

Functional work on Slo1 effectively describes the dual regulation of channel opening by voltage and calcium with nested MWC-type allosteric models^21–23^. Structurally distinct parts of the protein, the voltage-sensing domain (VSD) and cytosolic calcium regulatory domains (CTD), each couple to a membrane-embedded domain containing the ion permeation pathway (or pore-gate domain, PGD) and alter its conformational bias when triggered by their cognate stimuli (membrane voltage changes or intracellular calcium). It is implicit that percolation of channels through the landscape underlying gating involves several structural intermediates. Auxiliary subunits further modify this landscape, either by introducing additional intermediate states or by changing the transition energy barriers^24^, ultimately resulting in their unique functional effects on channel gating.

Structures of multiple orthologs of Slo1^25–28^ and its complex with the brain-specific β4 subunit^28^ have been determined in two key conditions. One of these captures Slo1 in the absence of divalent cations (the divalent-free DVF state), at presumed 0 mV, and the other in saturating concentrations of Ca^2+^, with a presumed Ca^2+^ ion occupying each of the two distinct categories of high affinity Ca^2+^-binding sites (the divalent bound DVB state)^29^. The structures have revealed vital details pertaining to the high conductance and calcium regulation of Slo1 channels. However, the structures of BK channels determined in the DVF states (for the functionally well characterized orthologs) have generally been of lower resolution, which limit their use in threading decades of biophysical observations onto molecular structures^30^. Furthermore, given the strong thermodynamic linkage between voltage and calcium regulatory pathways in Slo1^21,31,32^, it remains unclear what aspects of the structural changes observed thus far arise solely from voltage gating as opposed to calcium regulation. Deconvolving voltage and calcium dependent structural changes of Slo1 will be pivotal to not only obtain a more granular view of its allosteric gating, but they will also be crucial to understand how the auxiliary subunits affect gating transitions linked to one or both stimuli. For instance, the γ1 subunit is thought to profoundly affect voltage-gating with relatively modest effects on calcium regulation^14,33^ while β1 exerts robust effects on calcium-regulation^34–36^ but has relatively smaller effects on voltage-gating.

Towards this goal, in this study we pursued multiple high-resolution reconstructions of Slo1 under DVF conditions. First, using different mutant Slo1 channels which either stabilize a closed PGD or an activated VSD, we identify a key structural change in the VSD which we propose is critical for voltage-sensing. Second, we determine the structure of the hetero-octameric (4:4 stoichiometry) complex of Slo1 with γ1 (LRRC26) regulatory subunit. The architecture of the complex, together with extensive biochemical experiments, define key interactions and mechanisms that are central to the assembly of the complex. Guided by our structures and that of the previous DVB state, we hypothesize a mechanism that might underlie the functional effect of γ1 on the allosteric coupling of VSD activation to channel opening.

## RESULTS

### High resolution reconstructions of the hSlo1 channel mutants in presence of EDTA

For our structural studies we first targeted three Slo1 channel mutants which perturb its gating in three distinct ways (**Fig. 1A**). R207A neutralizes a positive charge on the exterior end of the S4 segment and shifts the conductance *vs* voltage (GV) curves of Slo1 leftward by >50 mV in the absence of intracellular calcium (**Fig. 1B and C**), although at 0 mV opening of R207A remains modest. This shift in gating is similar to that of other charge neutralizing mutations of this residue^37–39^ and likely arises from stabilization of an activated conformation of the VSD. A second mutation in the S6 segment, F315A, exerts a relatively modest effect on voltage-sensor activation^40^ but disfavors pore opening so profoundly that, at +100 mV, even in the presence of 300 μM intracellular Ca^2+^, mutant channels reached a P_O_ of only ∼0.01 (**Fig. 1D and E**) while wild-type Slo1 under such conditions reached a P_O_ of approaching 1. Given the two extremes of gating behavior exhibited by these two mutations, we anticipated that they may reveal structural differences that may occur while in DVF saline. The third mutant 2D2A/4D4A, neutralizes calcium binding to the RCK1 site (D362A, D367A) and the Ca^2+^ bowl (4D4A: ^894^DDDD^897^ to ^894^AAAA^897^) and abolishes high affinity calcium regulation of Slo1 while minimally perturbing voltage-dependent gating in absence of calcium^29^. These 3 mutants (in the background of hSlo1_EM_) were expressed in mammalian cells as eGFP fusion constructs, purified in digitonin micelles by a combination of affinity and size exclusion chromatography, and subjected to single particle cryoEM analysis, in the presence of 5 mM EDTA. Reconstructions of each of the three mutants revealed single high resolution classes which, upon refinement (with C4 symmetry), yielded maps with GS-FSC resolutions of 2.43 Å (for F315A), 2.6 Å (for 4D4A/2D2A) and 2.72 Å (for R207A) (**Supplementary Fig. 1** and **Fig. 1F**). Due to the high quality of our maps, we were able to build and refine accurate atomic models of the full-length channel (**Fig. 1G** and **Supplementary Fig. 2A**). At the backbone level, the structures of all three mutants superposed well with each other (**Supplementary Fig. 2A**) with backbone RMSDs of ∼1.0 Å. In our study we will refer to this quaternary configuration of the channel as the C state. Our models also match the earlier DVF state model of the hSlo1^28^ at the backbone level (RMSD 1.63 Å) (**Supplementary Fig. 2B**). However, we observed multiple non-protein densities in our reconstructions which provides mechanistic insights into the function of hSlo1.

**Figure 1.**
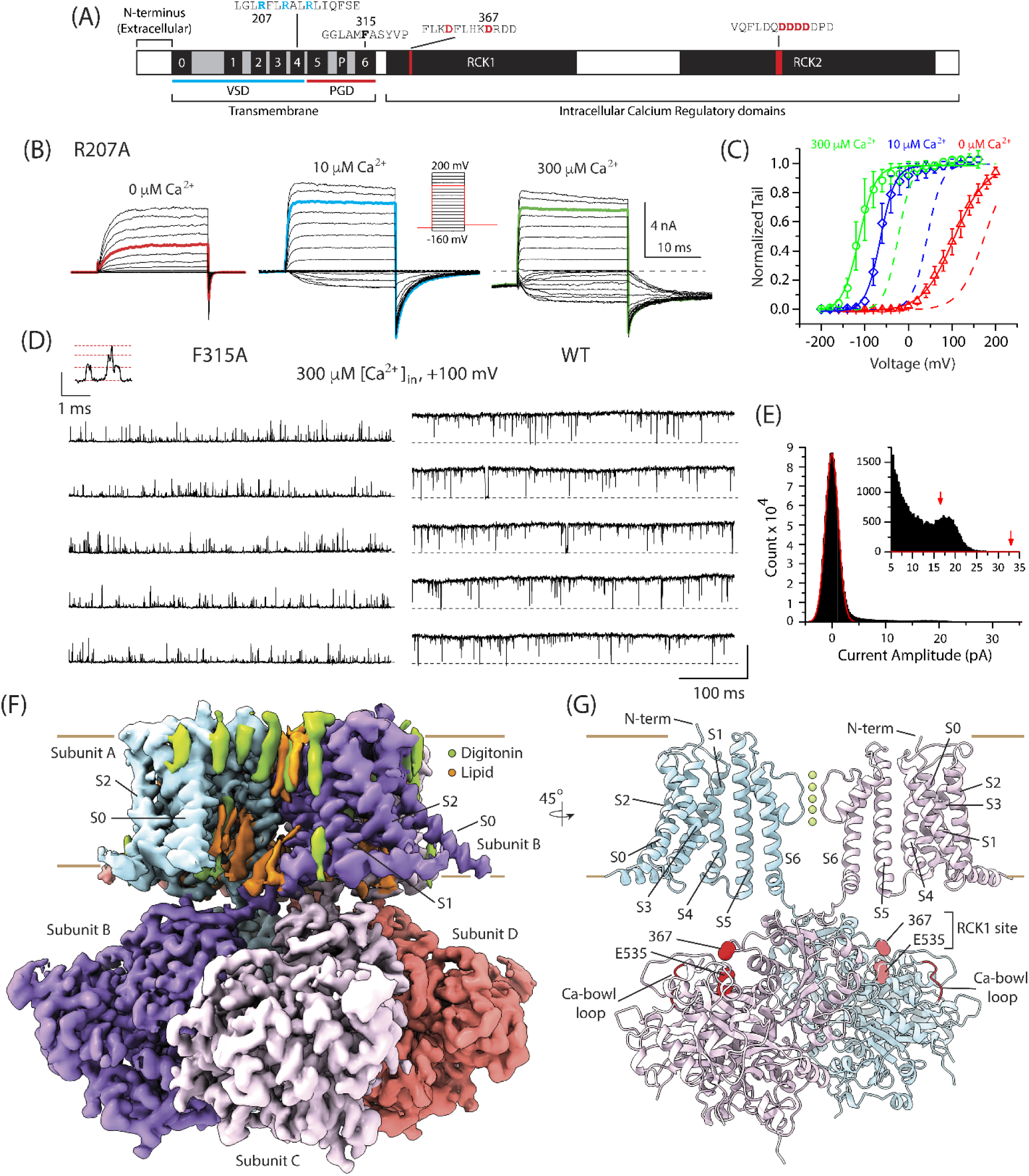
High resolution reconstruction of Slo1 channel mutants in EDTA. **(A)** Topological map of the Slo1 channel showing the organization of the transmembrane domain (with the voltage-sensing domain, VSD, comprising transmembrane helices S0-S4, and the pore gating domain, PGD, formed by transmembrane helices S5, S6 and the intervening P helix) and the large intracellular calcium regulatory domain (comprising the two RCK domains). The specific sites of the mutations explored in this study, and their adjoining sequence motifs, are highlighted. **(B)** Macroscopic currents of R207A evoked by a voltage protocol shown in inset in the presence of 0 (*left*), 10 (*middle*) and 300 (*right*) μM intracellular Ca^2+^. The dotted line marks the baseline (0) current level in these sample traces. **(C)** The G-V relationship of R207A determined from tail currents. Solid lines are Boltzmann fits to each GV: 104 mV, 0.70*e* (0 mM Ca^2+^), −64 mV, 1.49*e* (10 μM Ca^2+^), −114 mV, 1.33e (300 μM Ca^2+^). The dotted lines are the Boltzmann fit of WT BK channels: 174 mV, 0.88*e* (0 μM Ca^2+^), 43 mV, 1.47*e* (10 μM Ca^2+^), −23 mV, 1.60e (300 μM Ca^2+^). Error bars present standard deviations (n = 5) **(D)** Single channel activity of F315A (*left*) and WT Slo1 (*right*) recorded at +100 mV in 300 μM [Ca^2+^]_in_. Openings are upward with baseline marked by dotted lines. There were at least three channels in the representative recordings for F315A (*left*), as shown by three opening levels observed at +120 mV in the same patch (*inset*). The P_O_ of WT Slo1 in the representative recording (*right*) was 0.98 under identical conditions. Current scale represents 30 pA (main recordings and inset). **(E)** An all-point amplitude histogram generated from F315A single channel activity with a single Gaussian fit (solid red line) to the baseline. Most points not contributing to baseline, highlighted in the inset, represent unresolved openings and transitions and therefore do not result in well-defined amplitude components. Arrows highlight putative single channel levels assessed from a few longer dwells at open levels, suggesting that single channel conductance of F315A is markedly reduced compared to WT. P_O_ of this F315A channel was no more than 0.016 under the recording conditions. **(F)** Final unsharpened density map for the 2D2A/4D4A Slo1. The density for the four different subunits are colored uniquely. Non-protein densities corresponding to detergent (digitonin) and lipid (modeled as POPC) are highlighted in green and orange respectively. **(G)** Final model for 2D2A/4D4A derived from the map in (F) showing two diagonally apposed subunits (of the tetramer). All transmembrane domains and the two calcium binding loci in each subunit are marked. In this mutant, several of the aspartates in the acidic loop and D367 (together with D362) and mutated to Ala.

In all three reconstructions, several detergent and native lipid molecules were found, glued to the outer periphery of hSlo1 (**Fig. 1F**). Three lipid molecules are in positions suggesting they are of key significance. The first of these (outer pore lipid) is wedged in an inter-subunit groove formed by the external end of S6 of one subunit and the P helix of the neighboring subunit (**Fig. 2A**). A similarly localized native lipid has been identified in EM reconstructions of other Kv channels^41^ and might be stabilizing the outer pore. A second lipid molecule inserts its hydrophobic tail into an intra-subunit crevice (which we call the “medial crevice”) formed between the S5 and S6 helices (**Fig. 2A and B**). A third lipid guards a lateral fenestration between the S6 segments of two neighboring subunits, with its tail interacting with hydrophobic residues of the S6 segment and its headgroup tethered to a short stretch of positively charged residues (R329-K330-K331 or ‘RKK’ site) in the S6-RCK linker (**Fig. 2A and B**). In the divalent bound (DVB), open state of hSlo1^28^, the expansion of the S6 helices closes the medial crevice and the lateral fenestration. This requires the disengagement of these two lipid molecules from their respective binding sites. Hence both these lipid molecules preferentially bind and stabilize the C state of Slo1 and may regulate channel gating. Consistent with these inferences, MD simulations, complemented with functional experiments, have suggested that the interaction of the RKK site with phospholipid headgroups stabilizes the closed state of Slo1^42^. The lateral fenestration, a structural feature which has been observed in many homologous channels^43–45^, has also been suggested to form an access pathway for small molecules to enter the channel vestibule and regulate ion flux and channel gating. A similar mechanism is also possible in Slo1, but the guarding lipid would likely influence the accessibility and action of such pharmacological modulators. Additional detergent densities are also observed in the inner pore in many of our final density maps (**Supplementary Fig. 2D**).

**Figure 2.**
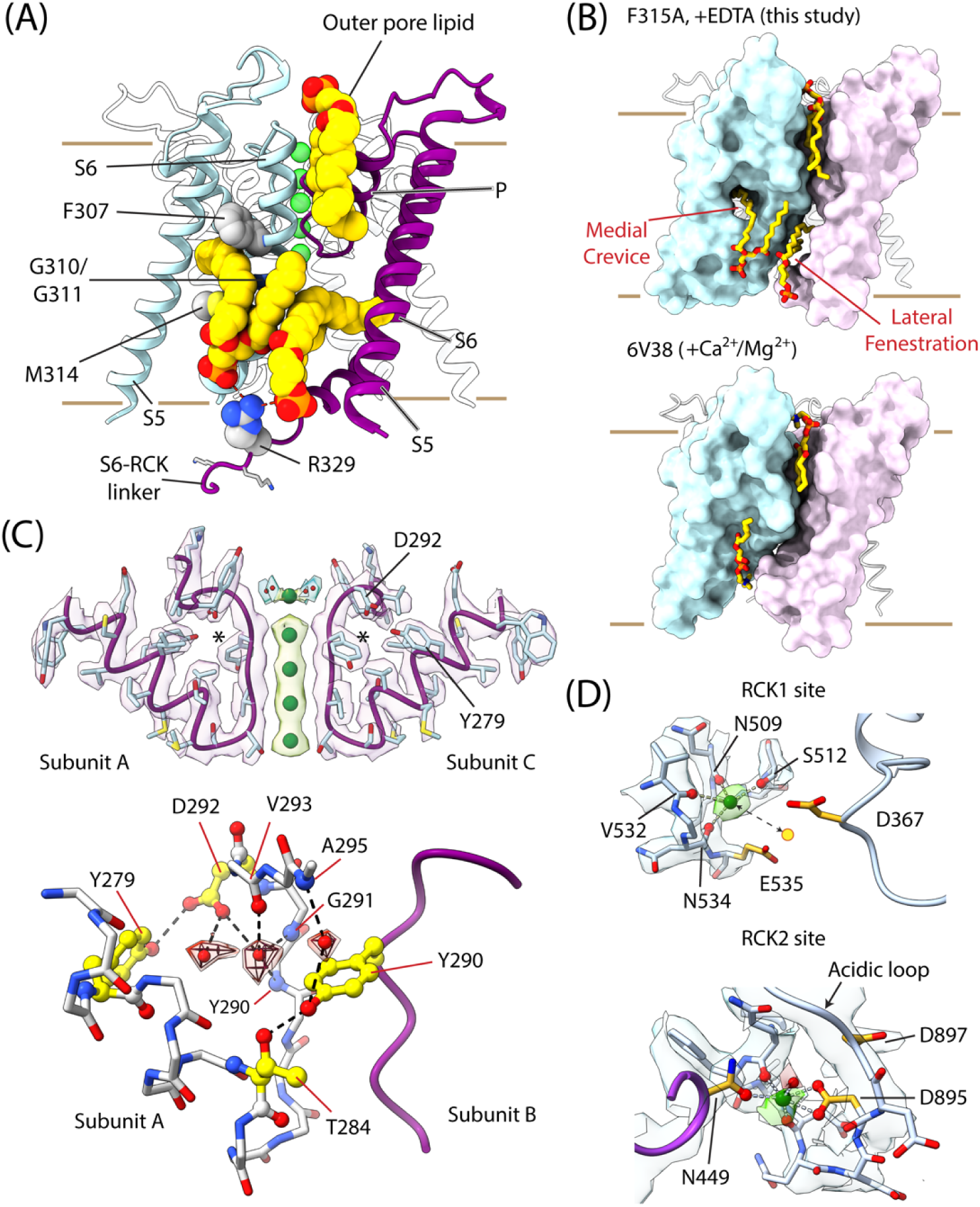
Lipid, water and ion interactions with Slo1 in presence of EDTA. All representations are derived from the model for the F315A mutant of Slo1. **(A)** The PGD showing the S5, S6 and P helices. Two adjacent subunits are colored in light blue and purple. The three key lipid molecules are shown in spheres (yellow: lipid tail; orange-red: phosphate head group). Sidechains of specific residues (F307, M314, R329) in intimate contact with the lipids are shown in spheres. Green spheres represent the K^+^ ions in the selectivity filter. **(B)** Surface representation of the PGD highlighting two adjacent subunits (in light blue and light purple) and the inter- and intra-subunit crevices that are lipid occupied in the F315A (+EDTA) model (*top*). The medial crevice and lateral fenestrations contract in the DVB state model (6V38)^28^ (*bottom*). **(C)** *Top,* Map and model of selectivity filter of two of the diagonally oriented subunits of Slo1 showing the K^+^ ion densities in the selectivity filter. 3 specific residues (D292, Y279 and Y290, marked with an *) are highlighted. *Bottom,* The selectivity filter of two adjacent subunits (one represented as sticks and the other purple ribbon), showing densities for three water molecules (superimposed mesh and surface: transparent red) and various protein (sidechain and backbone) interactions stabilizing them. All dashed lines represent distances < 3.2 Å. **(D)** Close-up views of the map and model around the RCK1 (*top*) and RCK2 (*bottom*) sites. *Top*, a tetrahedrally coordinated ion density (green, modeled as a K^+^ ion), is about 5.5 Å away from the mass center (yellow circle) of the sidechains of residues E535 and D367 (the high affinity Ca^2+^ coordinating sidechains in the RCK1 domain); *Bottom*, a density (in green), resembling a water-linked-ion, bridges the sidechains of N449 of one subunit and D897 of the adjacent subunit together with other backbone carbonyls in the acidic loop. All ion coordinations shown are < 3.2 Å.

Like other K^+^ channels, densities for 4 K^+^ ions are clearly visible (**Fig. 2C**) in the selectivity filter (S1-S4 sites) in all three of our reconstructions. A weaker density for a hydrated K^+^ ion at the S0 site is also observed. Behind the selectivity filter, we noted pseudo-spherical densities in the sharpened map for the F315A mutant (**Fig. 2C**). The positions of two of these densities closely match those of water molecules reported in the high-resolution crystal structures of prokaryotic K^+^ channels such as MthK and KcsA (**Supplementary Fig. 2E**). In KcsA these water molecules have been proposed to regulate inactivation gating associated with selectivity filter dynamics^46^. Although Slo1 channels are not known to intrinsically undergo such a process, mutations of side chains coordinating the water densities (D292, Y290 in the selectivity filter and Y279 in the P-helix) profoundly affect its ion conductance and gating^47–49^. Biochemically, size exclusion chromatography profiles of purified hSlo1 mutants, wherein these sites are perturbed by relatively mild substitutions (D292N and Y279F), show a robust destabilization of channel tetramers (**Supplementary Fig. 2F**). Furthermore, while in our preparative conditions, hSlo1 tetramers are comparably stable in low and high K^+^ (2.5 and 500 mM respectively) buffers, the two mutants show enhanced disassembly in low K^+^ buffers. Thus, this network of protein-water interactions regulates the stability and K^+^ ion interactions of the selectivity filter of hSlo1.

Close inspection of the high affinity Ca^2+^ binding sites in the F315A and R207A maps surprisingly showed clear ion densities (**Fig. 2D**), despite the presence of EDTA. The RCK1 density is tetrahedrally coordinated by backbone carbonyls of residues N509, S512, V532 and N534. It does not engage sidechains of D367 and E535, the principal determinants of Ca^2+^ binding at that locus^27,50^. An identically coordinated ion density at the RCK1 site is also seen in the 2D2A/4D4A map. Thus, it is unlikely to be a Ca^2+^ ion but could correspond to a K^+^ ion since it was the dominant cation in our buffers. The mean ion-O distances obtained from our refined models is 2.9 Å, matching the most frequent oxygen coordination distances for a K^+^ ion^51^. This ion is ∼5 Å away from the RCK1 Ca^2+^ coordination site. Through coulombic forces, it might influence the Ca^2+^ affinity as well as the divalent specificity of the RCK1 Ca^2+^ site. Larger divalents (such as Ba^2+^) would be repelled by the monovalent ion more strongly than smaller divalents (such as Ca^2+^ or Cd^2+^) perhaps explaining why RCK1 is more selective for the latter^52,53^.

The density in the Ca-bowl, while clear in the F315A and R207A maps (**Fig. 2D**), vanishes in the 2D2A/4D4A map. It is coordinated by the side chain of N449 of one subunit and, from the adjacent subunit, the backbone carbonyls and side chains of residues in the acidic loop (residues 892-900) which are known to be involved in Ca^2+^ binding. The acidic loop in the DVB state contracts around the Ca^2+^ density ^28^ but in our models for F315A and R207A it is somewhat more relaxed and adopts a significantly different configuration in the 2D2A/4D4A mutant (**Supplementary Fig. 2B**). Additionally, D897, which is one of the two most critical residues for Ca^2+^ binding at this site^54^, is twisted away from the Ca^2+^-bowl density in our F315A and R207A models. Thus, this density could represent a Ca^2+^ ion originating from contaminants in our buffers or inadvertently deposited by ash-fabricated filter papers used for plunge-freezing EM. Alternately, it could represent a K^+^ ion, which would suggest a competition between K^+^ and Ca^2+^ for this site. Although interactions with K^+^ will likely be of low affinity, in a cellular milieu K^+^ is orders of magnitude more abundant than Ca^2+^. Although more structural and functional investigations will be necessary to resolve this puzzle, we note that resolution of structures will be critical to resolve putative monovalent occupancies.

### Configuration of S4 charges in R207A, F315A and 2D2A/4D4A

Gating charge displacement *vs* voltage (QV) curve measurements have indicated that, at 0 mV, charge neutralizing mutants of R207 bias their VSDs towards their activated conformations^38^ while in F315A and 2D2A/4D4A the VSDs should prefer their resting conformations^32,40^. Yet at the backbone level the VSDs of all three mutants are superimposable (**Supplementary Fig. 2A**). However, a close inspection of the S4 charges reveals a striking difference (**Fig. 3A**). In F315A (and 2D2A/4D4A (**Supplementary Fig. 3a**)), the side chain of R207 is flipped upward, while that of R210 is featured right next to the conserved Phe residue in S2 (F160) and R213 is flipped downward. F160 forms the core of the hydrophobic gasket that intercepts the water accessible crevices within the VSD and focuses the electric field^55,56^. R210, being placed at the same level as F160, effectively seals this hydrophobic barrier and separates the interior from the exterior VSD crevices. However, in the VSD of R207A, the side chains of R210 and R213 each flip upward by ∼ 6 Å, such that R210 is housed in the external crevice and R213 moves out of the internal crevice and plugs the hole in the hydrophobic gasket. This suggests that the focused electric field is likely to be centered around the F160 position^38^ and the movement of R210 and R213 sidechains displace gating charges. Between the two charge configurations, the distances between the gating charges (R210 and R213) and negatively charged residues (particularly D186 and D153) lining the VSD crevice walls (in S1-S3) (**Fig. 3A**) change significantly. The changes in these interactions likely define the potential endpoints of the landscape of VSD activation. We propose that the VSDs of the mutants F315A and 2D2A/4D4A represent the resting state of hSlo1 VSD while that of R207A represents the activated state, respectively. VSD activation of Slo1 thus involves side-chain re-orientations of two S4 charges through a focused electric field and follows the “transporter model” of activation^57–59^. It is important to mention that in the previous DVF state models of Slo1^25,28^ the side chain orientations of the S4 charges were poorly constrained (**Supplementary Fig. 3B and C**) by the density maps due to their relatively low resolution, while in our study the high resolution maps enable us to make these precise structural inferences.

**Figure 3.**
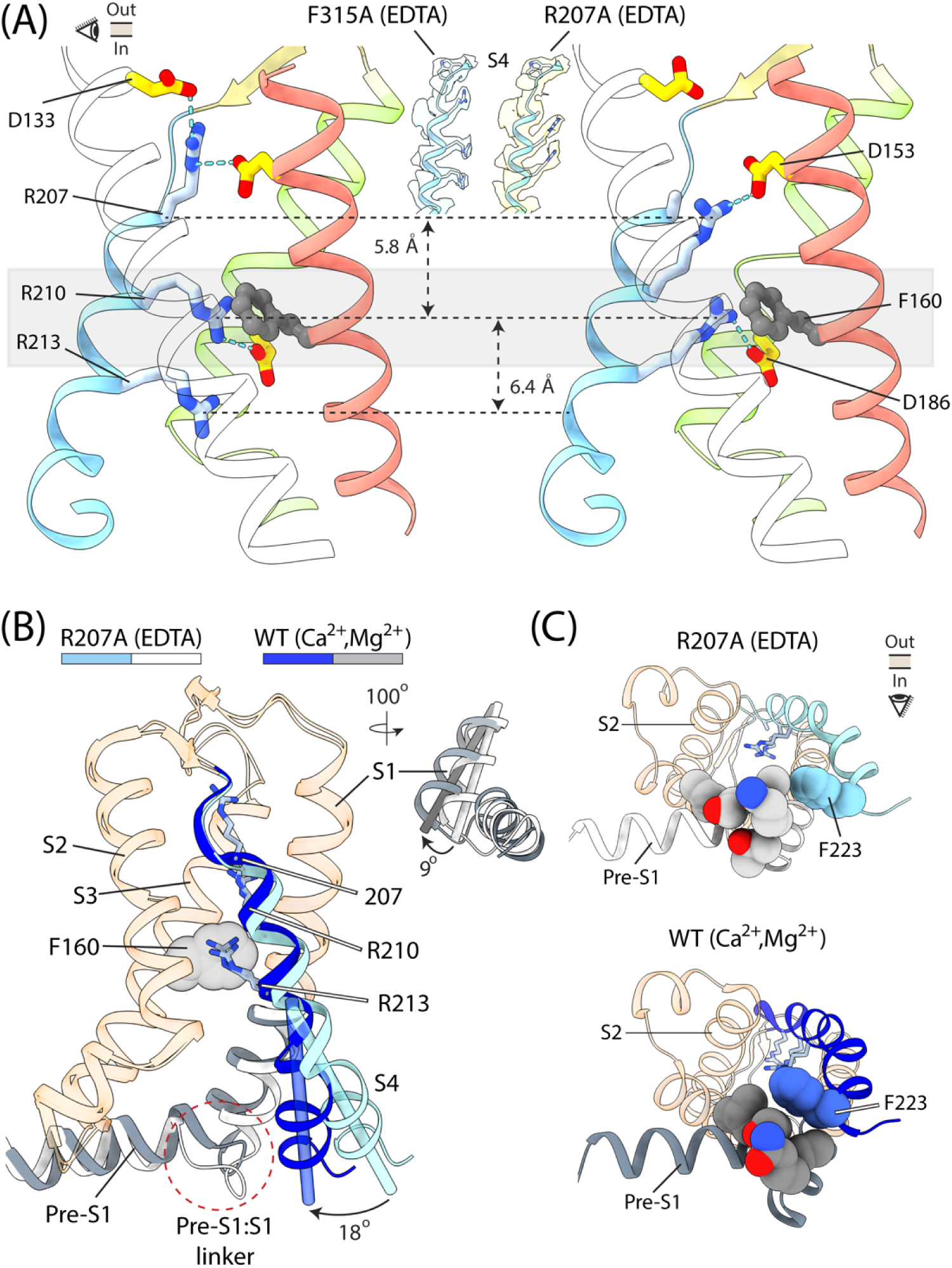
Re-arrangements in the VSD of Slo1. **(A)** VSDs of F315A (*left*) and R207A (*right*) in presence of EDTA (S1: transparent white, S2: red, S3: green and S4: blue; S0 is not shown) showing sidechains of R210, R213 and R207 (*left*) or A207 (*right*). The focused electric field is speculatively marked by the transparent gray slab. Blue dotted lines indicate contacts < 3.5 Å. Between the two states, D186 and R210 change their salt-bridge partners. Insets show the density of S4 charges in the resting (F315A) and activated (R207A) states. **(B)** Superposition of the VSDs of the R207A model and the DVB state model (6V38)^28^ (ref). S4 helix is in light blue/dark blue for R207A/DVB state models. The pre-S1 and intracellular end of S1 are in white/gray for R207A/DVB. The inner end of S4 and S1 rotate by 18° and 9° respectively. Other VSD helices align well between the two models (transparent red helices). **(C)** Bottom-up views of the VSD of R207A (*top*) and DVB state (*bottom*). F223 becomes more effectively packed against residues connecting the pre-S1 helix to the inner end of S1.

Comparison of the activated VSD of R207A with that of VSD of the DVB state^28^ (ref) shows that the gating charges (R210 and R213) are oriented similarly (**Fig. 3B**). However, in the latter, the C-terminal, intracellular end of S4 bends inward by ∼18°, towards the internal end of S1. This transition likely occurs after gating charge movement. In concert with or following this, the S4-S5 linker moves facilitating the expansion of S6 helices. Between the activated VSDs of R207A and DVB states, the intracellular end of S1 also rotates by ∼9° and the short linker connecting S1 to the pre-S1 helix also readjusts. As a result of these changes, F223 at the intracellular end of S4, becomes more effectively packed against the pre-S1:S1 linker in the DVB state relative to the C state of R207A (**Fig. 3C**). F233 has been suggested as a critical determinant of electromechanical coupling in Slo1^60^. Several perturbations in the C-terminal end of S4 have also been proposed to compromise allosteric coupling between the VSD and the pore^37,38,61^. Additionally, Mg^2+^ binding has been shown to favor channel opening by strengthening electromechanical coupling^62–65^. While the pre-S1 helix might ordinarily be dynamic, Mg^2+^ binding (regulated by D99 in the preS1 helix, N172 in intracellular loop linking S2-S3 helices and E374/E399 on the RCK1 N-lobe) likely makes it more rigid and effectively engage the intracellular end of S4. Together, these structure-function correlations hint towards an important role of the inner ends of VSD helices in facilitating expansion of the inner ends of the S6 helix, in the later steps of voltage gating.

### Architecture of the Slo1-LRRC26 complex

Voltage-dependent activation of Slo1 is dramatically facilitated by the LRRC26 (or γ1) subunits^14^ (**Fig. 4A** and **Supplementary Fig. 5A,B**) and we wondered whether γ1 might influence VSD status in DVF conditions. To understand the structural principles underlying Slo1-γ1 assembly and the mechanisms by which γ1 modulates Slo1 gating, we performed single particle reconstructions of hSlo1:LRRC26 complex in presence of 5 mM EDTA (**Supplementary Fig. 4A**). We initially obtained a reconstruction of the complex where the extracellular density for the LRR domains was relatively weak (**Fig. 4B**). The corresponding map was further refined by masking out the extracellular Leucine Rich Repeat Domain (or LRRDs), to obtain a 3.1 Å reconstruction of the complex, where densities for γ1 transmembrane segments were clearly visible (**Fig. 4A** and **Supplementary Fig. 5C**). In an alternate workflow, the density for the intracellular gating ring was masked out and reconstruction analyses resulted in a single C4 symmetric class of the 4:4 complex of hSlo1_EM_-γ1 at a GS-FSC resolution of 3.13 Å where the LRRD and TM (transmembrane segment) of γ1 were clearly resolved (**Fig. 4C** and **Supplementary Fig. 5C**).

**Figure 4.**
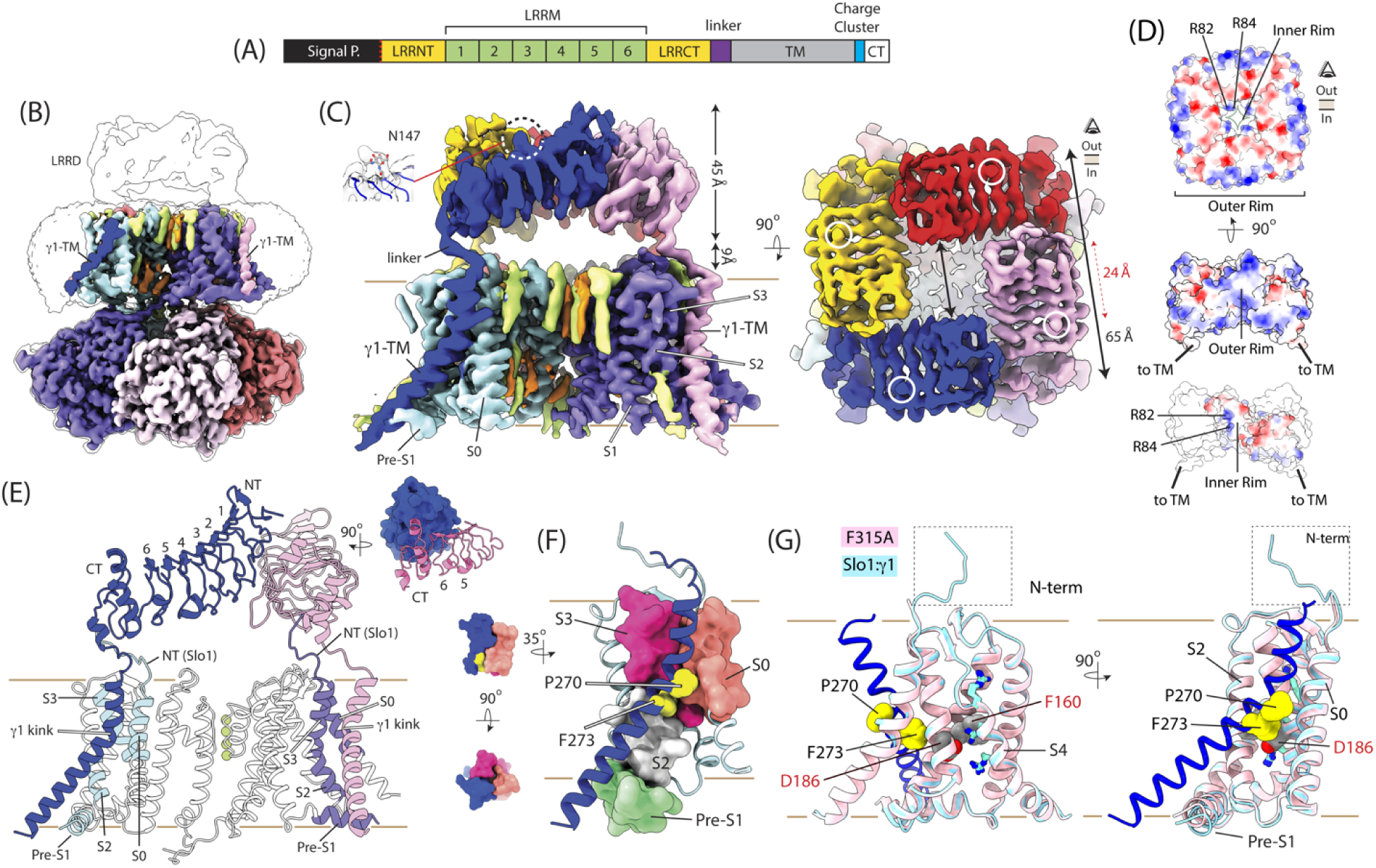
Structure of Slo1:γ1 complex in EDTA. **(A)** *Top*, Topological map of the γ1 subunit showing the key structural motifs. The N-terminal signal peptide is cleaved post-translationally. **(B)** LRRD-masked unsharpened density map (colored), contoured at a high threshold, is superposed on the unmasked unsharpened density map, contoured at a low threshold (transparent white). The densities for γ1-TM of 2 subunits are indicated in the LRRD-masked map. **(C)** *Left,* Side view of the Gating-Ring masked (GR-masked) unsharpened map for the Slo1:γ1 complex where different subunits are colored uniquely. Detergent and lipid densities are in green and orange, respectively. The inset shows glycosylation density on residue N147. *Right*, Top view of the GR-masked unsharpened map showing the LRRD density and highlighting the glycosylation density corresponding to N147. **(D)** Surface electrostatic potential maps of the tetrameric LRRD ring in a top-down view (*top*) and side view (*middle*) (Blue: 10kT/e, White: 0 and Red: −10kT/e). *Bottom*, electrostatic potential of the inner rim of a single LRRD (only 3 LRRDs are shown, with 2 LRRDs in white and only 1 colored). **(E)** Model of the transmembrane domains of two adjacent subunits of Slo1 and their γ1 partners. The various motifs of the LRRD and the kink in the γ1-TM are marked. Specific segments of the VSD which pack against the γ1-TM are colored (in green in one subunit and purple in the other). The inset shows the interface between the LRRDs of two adjacent γ1 subunits. **(F)** Parts of the S0, S2, S3 and pre-S1 helix are shown in surface representation, together with the γ1-TM. The latter (in particular two residues, P270 and F273 shown in spheres) fits into surface grooves formed by specific regions of the Slo1 VSD. **(G)** Superposition of the VSD of the F315A model (resting state), in pink, with that of the Slo1:γ1 complex (GR-masked), in light blue. The γ1-TM is in dark blue. Key residues of the VSD (S4 charges, F160 and D186) are perfectly aligned between the two models. F160 and D186 (labeled in red) point within the VSD core and the kink of γ1-TM (particularly F273) is featured right next to them.

In the quaternary complex, the LRRDs of the four γ1 subunits are organized as a ring of “interlocked dominos” (**Fig. 4C**). The ring is 45 Å thick along the membrane normal and lies ∼ 9 Å above the membrane. Along the membrane plane, the external edge (or outer rim) of the LRRD ring is ∼ 65 Å long. A central gap (∼24 Å in lateral dimension) in the LRRD layer directly connects the external milieu to the channel pore. The LRRD ring features modest surface electrostatic characteristics (**Fig. 4D**). Its outer surface (parallel to the membrane), directed away from the channel, is somewhat electronegative while the outer and inner rims (perpendicular to the membrane) are slightly electropositive. Each LRRD is shaped like a curved solenoid comprising six Leucine Rich Repeat Motifs or LRRMs 1-6, each forming a β-turn-α loop (**Fig. 4E**). Two additional LRRMs, LRRM-NT and LRRM-CT, flank the core LRRMs 1-6 on the N and C terminal ends, respectively. LRRM-NT and LRRM1-2 of one subunit form an interface with LRRM-CT and LRRM6 of the neighboring subunit burying ∼550 Å^2^ of molecular surface between them (**Fig. 4E**). The LRRDs do not contact the Slo1 channel, except for the LRRM-CT which interacts with the extracellular N-terminus of Slo1 (**Fig. 4E and F**), consistent with the results of LRET measurements^66^. A structural consequence of this interaction is that 7-8 residues of the Slo1 N-terminus, which were invisible in the reconstructions of the Slo1 mutants, become ordered and visible in the γ1 complex. We also note a clear bump at N147 in our map (**Fig. 4C**), likely corresponding to glycosylation. N147 is on the external surface of LRRD, directed away from the channel and the membrane. In the absence of glycosylation of γ1, gating shifts are not observed^67^, but it is not known whether this is a lack of function or assembly.

The transmembrane helix of the γ1 subunit interacts intimately with the VSD of Slo1. It is kinked on the extracellular side around residues 269-273 (**Fig. 4F** and **Supplementary Fig. 5D**). Above the kink, the helix fits into a groove formed by the extracellular ends of the S0 and S3 helices of Slo1 (**Fig. 4F** and **Supplementary Fig. 5E**). Below the kink, the γ1 transmembrane helix leans against the S2 and pre-S1 helices. The transmembrane helices of hSlo1 in this complex are identical to that in the C state, with the voltage-sensing R210 and R213 rotamers in a downward flipped state, representing a resting VSD (**Fig. 4F**). Interestingly, the kink in the γ1-TM is positioned intimately next to Slo1 residues (for example, F160 and D186) that are critical for its voltage dependent activation (**Fig. 4G** and **Supplementary Fig. 5E**). The quaternary arrangement of gating ring, as inferred from the LRRD masked map, matches that of the C state (for the Slo1 mutants) (**Supplementary Fig. 5C**). Density corresponding to the C-terminal end of the γ1 subunit (residues 294-330) was not visible in our maps indicating that in this conformational state it is likely to be relatively flexible.

### γ1-TM kink is critical for assembly and regulation of Slo1

Co-expression of hSlo1EM and γ1 as eGFP and mCherry fusion constructs allowed us to use Dual Color Fluorescence Size Exclusion Chromatography (DC-FSEC)^68,69^ together with a two-step affinity purification process to probe the regions of γ1 that are central for its assembly with Slo1 (**SupplementaryFig. 6A-C, Fig. 5A and B**). Under our expression and purification conditions about ∼50% of total Slo1 is in complex with γ1. In comparison, >90% of total Slo1 forms a complex with β4 and β2 subunits (**Fig. 5B**). The efficiency of assembly of the four homologous γ subunits followed the rank order γ1 > γ2 > γ3 ∼ γ4. For γ3 and γ4, hSlo1_EM_-γ complexes comprised <10% of total Slo1 expressed. We also compared the assembly efficiencies of two chimeric γ subunits. The chimera, γ13 (LRRD from γ1; TM and C-terminus from γ3) assembled poorly with hSlo1_EM_ (at levels similar to WT γ3) while γ31 (LRRD from γ3; TM and C-terminus from γ1) assembled much more efficiently (almost like γ1) (**Fig. 5B**). Thus the γ-TM segment strongly favors formation of the Slo1-γ complex.

**Figure 5.**
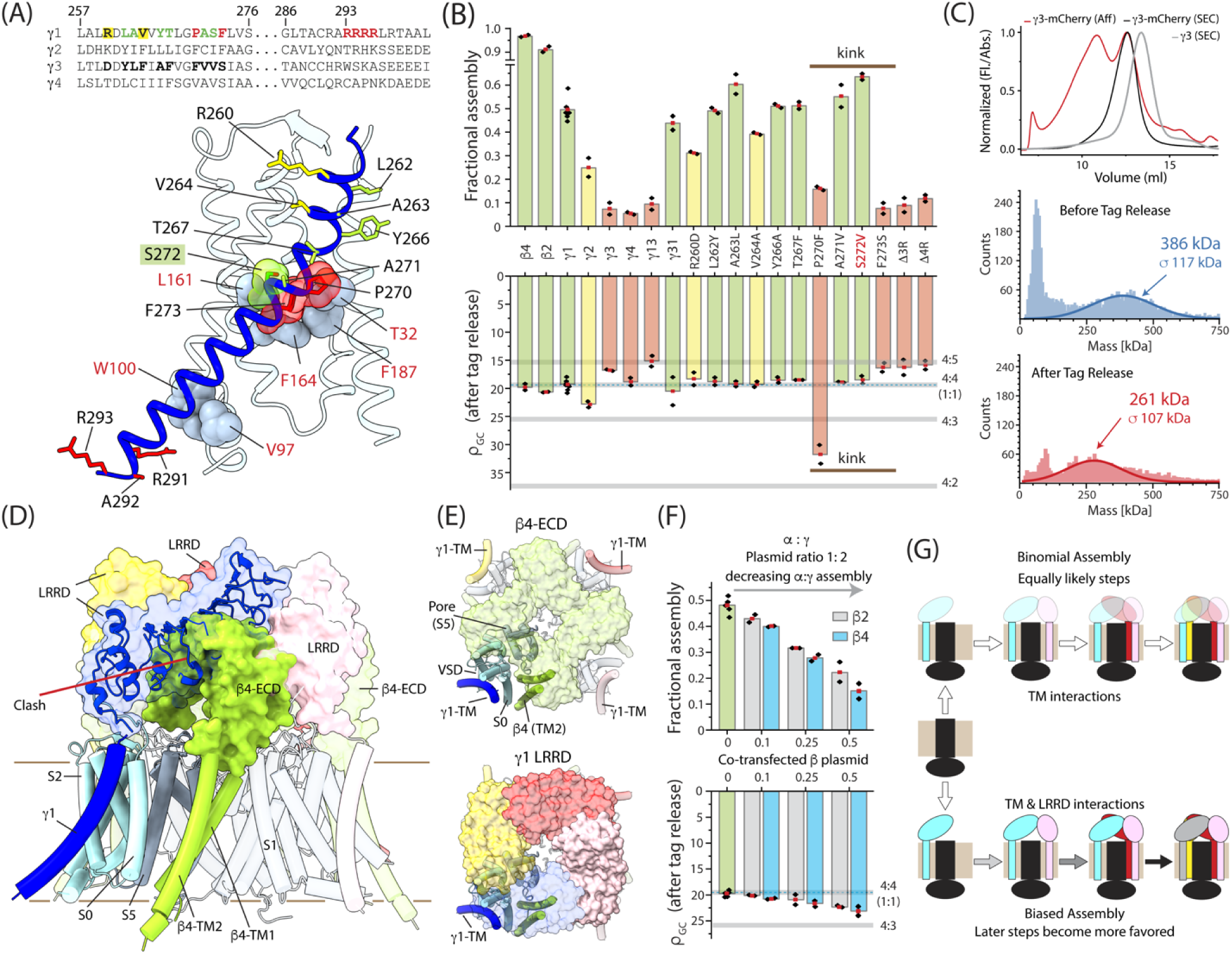
The transmembrane helix and LRRD of γ1 influences its assembly with Slo1. **(A)** *Top,* Sequence alignment of the transmembrane helices of γ1-4. Specific residues of γ1 tested for their impact on assembly are marked in bold and colored as: green - little or no reduction; yellow: medium reduction; red: large reduction. Corresponding residues in γ3 are also marked. *Bottom*, Structure of the γ1-TM and the Slo1 VSD, showing sidechains of γ1 residues tested (marked and colored as in the sequence alignment). S272, P270 and F273 (all at the γ1-TM kink) are highlighted in spheres. Slo1 residues which putatively interact in γ1-TM are represented as grey spheres (red labels). **(B)** *Top*, Fractional assembly of different auxiliary subunits (and their mutants) with Slo1 measured as fraction of Slo1 in complex with auxiliary subunits to total Slo1. *Bottom*, ρ_GC_ (ratio of the eGFP:mCherry fluorescence) obtained after tag release from doubly purified Slo1:auxiliary subunit complex. Horizontal lines indicate the calibration values of ρ_GC_ obtained from a control dimeric membrane protein (for 1:1 equivalent to 4:4 stoichiometry) and from mixtures of free eGFP/mCherry (for 4:5, 4:4, 4:3 and 4:2 stoichiometries). **(C)** *Top*, Size exclusion chromatography profile of affinity purified full-length γ3-mCherry (monitored with mCherry Fluorescence) in 0.01% L-MNG buffer (red). Fractions corresponding to the smaller molecular weight peak were pooled and exchanged into detergent free buffer by size exclusion chromatography (before and after removal of the mCherry tag) where the elution was monitored via A280 absorbance (black and gray traces). *Middle, Bottom,* Mass photometry profiles of the lower molecular species in affinity purified γ3-mCherry, before (*middle*) and after (*bottom*) release of the mCherry tags. **(D)** A composite model of the Slo1:γ1:β4 complex generated by Slo1 guided alignment of our structural model of the Slo1:γ1 complex (GR-masked) and the structural model of the Slo1:β4 complex. One β4 subunit is shown in opaque green (other β4 subunits are in transparent green). One γ1-TM is shown in opaque dark blue (others are in transparent pink, red and yellow). One Slo1 subunit is opaque with the VSD helices in light blue and the PGD in gray. The LRRDs of the 4 γ1 subunits and the extracellular loop of the β4 are shown as surface representation (LRRD of 1 γ1 subunit also shown in cartoon). The clash between the extracellular domains (ECDs) of the β and γ subunits in this hypothetical dodecameric ternary complex is indicated. **(E)** Top down views of the ECDs of β4 (*top*) and γ1 (*bottom*) of the hypothetical dodecameric ternary complex. Transmembrane helixes of 1 subunit each of Slo1, β4 and γ1 are shown. **(F)** Effect of co-expression of β4 and β2 subunits on fractional assembly of Slo1 with γ1 (*top*) and ρ_GC_ of doubly purified Slo1:γ1 complex at different levels of β subunit transfection. **(G)** A non-binomial, cooperative model of assembly of Slo1 tetramer with γ1 where interaction between LRRDs makes high stoichiometry complexes more favorable (*lower path*) as opposed to a binomial, independent model of assembly (*upper path*) where the assembly of γ1 with Slo1 tetramer does not depend on the number of γ1 subunits pre-associated with the Slo1. Increasingly darker arrows in the steps of association (in the cooperative assembly model) indicate an increase in avidity.

Additional mutants that were tested replaced specific residues of γ1-TM with corresponding residues in γ3 (**Fig. 5A**). Most of them minimally or modestly perturbed assembly with Slo1 (**Supplementary Fig. 6D**). The clear exceptions were P270F and F273S which dramatically reduced complex formation (**Fig. 5B**). Both residues are located at the kink of the transmembrane segment. The side chain of γ1 F273 occupies a hydrophobic pocket formed by Slo1 channel residues, L161, F164 and F187 (on S2 and S3), and γ1 P270 is close to the sidechain of T32 (on S0) possibly connected via a hydrogen bond (**Fig. 5A**). These two γ1 mutants likely destabilize the kink or disrupt the local intimate packing with Slo1 and highlight the structural importance of the kink. Two additional mutants targeted the C-terminal end of the γ1-TM, deleting multiple positively charged residues (**Fig. 5A**). These mutants also robustly compromised the ability of γ1 to associate with Slo1 (**Fig. 5B** and **Supplementary Fig. 6D**). This region of γ1 comes close to the pre-S1 helix of Slo1 (**Fig. 4D**) but due to the limited density in this structural region, we are unable to ascertain specific structural contacts. Positively charged residues at the ends of a TM facilitate its membrane integration due to interaction with phospholipid head groups^70,71^. Thus, it is possible that the charge cluster of γ1 affects its association with Slo1, by influencing the insertion or orientation of the γ1-TM in membranes. Some γ1 mutants studied here produce much smaller shifts in Slo1 GVs relative to WT-γ1^72,73^. It has been unclear to what extent the loss of function arises from disruption of allosteric linkage between γ1 and Slo1 or the inhibition of complex formation. This is reminiscent of the “binding-gating conundrum” for ligand gated channels^74^, where a mutation can thwart ligand dependent channel opening by inhibiting ligand binding or severing allosteric linkage between ligand binding and downstream conformational changes. The S272V mutation (also featured at the γ1-TM kink) is an exception. Our experiments show that it assembles with Slo1 as efficiently as (if not slightly better than) WT-γ1 while previous functional results have shown that it substantially reduces shifts in Slo1 GVs relative to WT-γ1^72^. Hence, an intact γ1-TM kink is critical both for γ1 assembly and its modulatory effects.

### Role of LRRDs in complex assembly

The ratio of the eGFP:mCherry fluorescence intensity (ρ_GC_) of the doubly affinity purified Slo1-γ1 complexes allowed us to infer their average stoichiometries (see Methods). The specific value of ρ_GC_ was calibrated to defined stoichiometries using a control dimeric protein and purified free mCherry and eGFP (see Methods). Based on this, we found that under our experimental conditions, the average stoichiometry of purified Slo1-γ1, Slo1-β2 and Slo1-β4 complexes was 4:4 (**Fig. 5B**). Across all γ variants tested in this study, the inferred stoichiometry was similar, except for P270F, where it was ∼4:2.4 indicating that under specific conditions Slo1-γ1 can form complexes with fewer than 4 γ1 subunits.

In many of the γ1 mutants, which poorly assembled with Slo1 (Slo1 in complex was <∼ 10% of total Slo1), the stoichiometry of the purified complexes were inferred to be 4 Slo1: > 4 γ. This is unexpected and could potentially reflect improperly assembled complexes. But why is there such a preference for high stoichiometric states? Slo1-γ1 complexes have been proposed to assemble into four α:γ1 stoichiometric combinations^9,75,76^. If all the combinatorial assemblies were equally likely or if a single copy of the γ1 subunit assembled with Slo1 tetramers independent of another, then we would’ve expected the average stoichiometry to decrease (4 Slo1: < 4 γ) in mutants where the efficiency of assembly is reduced. One possible structural explanation underlying this stoichiometric bias could be that the LRRDs interact with each other (as seen in the structure) and this favors complexes with two or more γ subunits. Such a hypothesis would demand that the LRRDs have an intrinsic tendency to interact with each other, possibly even in absence of Slo1. To test this, γ1-4 subunits were expressed individually as mCherry fusions without Slo1, affinity purified in L-MNG micelles, and examined via FSEC (**Fig. 5C** and **Supplementary Fig. 5H**). A broad peak corresponding to polydisperse, high order oligomer(s) was observed in all four cases. For γ2-4, a second relatively sharper peak corresponding to a lower order oligomer was also seen. For γ3, the species corresponding to smaller oligomeric peak was further analyzed by mass photometry^77^ (**Fig. 5C**). The mass of this protein-detergent complex, measured before and after release of the mCherry tag, indicated a net reduction of 125 kDa due to tag release. Considering each tag is ∼29 kDa, the result indicates that the corresponding γ3 species most likely corresponds to a tetramer or a mixture of tetramer and pentamer. The LRRDs of γ1-4 (TM and C-terminal ends deleted), when expressed and purified, are polydisperse showing various degrees of oligomerization (**Supplementary Fig. 5I**). Both experiments support intrinsic homomeric interaction between the LRRDs of γ1-4. These experiments, however, do not clarify if the self-interactions between the LRRDs are important for γ1 assembly with Slo1 tetramers.

The structure of the Slo1:β4 complex shows that the 2 TM segments of β4 associate with the channel, each at a different location than the γ1-TM (**Fig. 5D**)^28^. This raises the possibility that γ1-TM and β4-TMs could be accommodated in the same complex. Previous functional work has provided two differing answers to this question. Whereas in HEK or LNCaP cells co-expression of β1 subunit substantially reduced or eliminated the γ1-mediated shift effect^14,78^, it was also observed that β2 and γ1 can both mediate subunit specific functional effects in single BK channels when co-expressed in oocytes^79^. We therefore asked whether a ternary complex containing the large extracellular loop between the 2 TMs of the β subunits would structurally clash with and possibly disassemble the ring of LRRDs. To test this, we co-transfected Slo1-eGFP with γ1-mCherry (plasmid weight transfection ratio of 1:2) together with different amounts of β4, with a His tag on its C-terminus. Following the same biochemical strategy as before, we observed that the fraction of Slo1 in complex with γ1 decreased with increasing levels of co-transfected β4 cDNA (**Fig. 5F** and **Supplementary Fig. 6E**). Interestingly ρ_GC_ in the purified complexes slightly increased with increasing amount of transfected β subunits suggesting that in these complexes tetrameric Slo1 possibly assembles with fewer than 4 γ1 subunits. At the highest amount of β4 tested, the inferred average stoichiometry was 4 Slo1: 3.5 γ1. Similar results were obtained with β2. We were not able to confirm whether Slo1, β2/4 and γ1 formed a ternary complex because of non-specific interactions of Slo1 or γ1 with the His-tag (on β subunit) affinity capture resin (but inspection of the FSEC profiles of the γ1-free and γ1-contaning complexes suggest that the β- and γ-subunits might prefer to segregate into different complexes **Supplementary Fig. 6F and G**). The implication of these results is that although the TMs of γ1 and β interact with Slo1 at non-interfering interfaces, the clash between the LRRD ring and the extracellular loop impedes their simultaneous association. This suggests that, if the structural interaction between the LRRDs is disrupted, it impedes the ability of γ1 to associate with Slo1. Consistent with this, deletion of the LRRD segments or replacing it with unrelated globular domains in the context of γ1 alters the functional effects of mutant subunits^67^ probably due to aberrant assembly. Designer constructs, in which the γ1-TM replaces the second transmembrane helix of β subunits, are however able to efficiently assemble with and modulate Slo1, even without the LRRD^67,75^. Thus, while the LRRDs are important for the association of native γ subunits with Slo1, they are likely vestigial for their modulatory functions. Furthermore, the role of LRRDs in chaperoning the Slo1: γ1-TM union can be fulfilled by other engineered protein modules. Overall, we propose a model for Slo1:γ1 assembly in which the initial association of a single γ subunit with a Slo1 tetramer is driven by the interaction of its TM with the non-S4 helices of Slo1-VSD. Thereafter, the association of the second γ subunit becomes more favorable due to the interaction between the two LRRDs (**Fig. 5G**) and this possibly underlies the bias towards the high stoichiometric states. Thus, the association of the multiple copies of γ1 subunit with Slo1 is not independent and likely follows a non-binomial, cooperative model. Our results and model do not exclude the possibility that channels with less than 4 γ1 subunits may form for instance under conditions of very low γ1 expression^9,75,76,79^.

## DISCUSSION

Over the last several decades substantive efforts, combining electrophysiological, spectroscopic^80–82^, and chemical-biological^83,84^ methods, have been invested to decipher how Slo1 VSDs sense membrane voltage and regulate channel opening and how this process is influenced by the quaternary gating ring. The structures of Slo1 reported until now have provided a foundation to start connecting many of these dots. Yet due to the limited resolution of the single resolved conformation of Slo1 under DVF conditions, central questions regarding voltage-gating remained unanswered.

### VSD activation in Slo1 channels

Here, we determined high resolution structures of human Slo1 mutants which dramatically change the conformational bias of Slo1 (at 0 mV and nominally calcium free conditions) to VSD-resting/Pore-closed or VSD-active/Pore-closed states. Comparison of these structures show that rotameric flipping of key S4 arginines (R210 and R213) displaces their positively charged guanidium moieties by ∼6 Å through a focused electric field without virtually any discernable change in the S4 helix. These changes are reminiscent of the voltage-sensing mechanism proposed for the voltage-sensing phosphatase^85,86^, where a small vertical displacement of the S4 together with rotameric flips of S4 arginines were observed between structures of a Fab-stabilized resting VSD and a mutant stabilizing the active VSD.

This small sidechain movement in Slo1 might be sufficient to account for the ultra-fast gating currents observed in the functionally well-characterized orthologs of Slo1^87^ (refs). Our observed changes would also be consistent with the emergence of the omega- or gating-pore currents in a charge neutralized Slo1 mutant (R210H)^38^. Since R210 plugs the hydrophobic gasket in the resting state, ions are prevented from passing through a resting VSD, possibly via coulombic repulsion. In contrast, when R210 is neutralized in R210H, this might enable cations to pass through when the VSD is resting as a consequence of an altered electrostatic environment around the 210 site. Upon VSD activation, R213 then plugs the gasket and turns off ion flux. These structural inferences line up with the transporter model of voltage-sensor activation^57,58,88^. While the latter model was originally proposed in the context of Shaker K_V_ channels, direct structural evidence of this proposal had been lacking until structures of the TPC1 channel revealed multiple conformations of their VSDs^89^. These defined a probable sequence of transitions in the VSDs, mimicking the conformational transitions of transporters accompanying substrate transport. Our structural observations with Slo1 reinforce this hypothesis and contrasts Slo1 from other voltage-gated channels where the dynamics of movement of S4 helices can be substantially larger^90^.

Our results also provide information pertinent to the identities of voltage-sensing residues in Slo1. While a recent study argued that R210 and R213 are the primary gating charges^38^, an earlier study reported that R213 (but not R210) and negatively charged residues in S2/S3 segments^37^ contribute to gating charge movement. While our results appear to be more consistent with the first hypothesis, the boundaries of the focused electric field would need to be properly defined to infer the contribution of the specific charge movement associated with the rotamer flip at each position to the total gating charge.

### The assembly of Slo1:LRRC26 complexes

To stabilize and thereby structurally trap conformational intermediates during voltage-activation Slo1, we determined the Ca^2+^-free structure of the hSlo1:hLRRC26 (γ1) complex which open at much lower depolarizations than WT hSlo1. Under our conditions of reconstruction, γ1 appears to have little effect on the structure of Slo1 except for a modest change in the position of the pre-S1 helix (**Supplementary Fig. 5C**). Nevertheless, our structural and biochemical experiments revealed two key principles involving Slo1: γ1. First, the single TM of γ1 features a kink that is possibly stabilized by intramolecular H-bonds and is intimately packed against non-S4 helices of the VSD. Under our experimental conditions, mutations at this locus (P270 and F273) dramatically affect the efficiency of γ1 to assemble with Slo1. Consistent with this idea, two γ1 paralogs (γ3 and γ4) which natively feature dramatic substitutions at this locus, assemble very poorly with Slo1. We note that, under heterologous expression conditions, the rank order of assembly of the four γ paralogs parallel the order of shifts in Slo1 GVs observed when they are individually co-expressed with Slo1^20^. It remains somewhat unclear to what extent the differences in these functional effects may arise from differences in assembly efficiencies or modulatory potencies. The impact of modified assembly must be appropriately considered in future mechanistic explorations of Slo1 regulation by γ1 via mutational analyses. A second important aspect of the Slo1: γ1 assembly that we discovered is the impact of LRRDs. These extracellular domains form a ring of interlaced dominos on top of the channel and the interactions between these domains bias Slo1: γ1 complexes towards high stoichiometric states. We hypothesize, therefore, that γ1 assembles with Slo1 cooperatively such that under heterologous expression conditions there would be a preponderance of channels in the 4:4 or 4:3 stoichiometric states. Under specific situations, however, low stoichiometric complexes (fewer than 4 γ subunits) may also form (such as in the case of the P270F mutant or upon co-expression of β subunits). To date, the best direct functional evidence in support of the formation of such “sub-stoichiometric” complexes comes from the exploration of Slo1 modulation by a β2-γ1 chimeric construct^75^, which lacks the LRRD.

The importance of the LRRD interactions in the overall complex assembly is further highlighted by an overall decrease in isolated Slo1-γ1 complexes in the presence of β subunits. Our analysis suggests that this might arise from a structural clash between the extracellular domains of β and γ. While our biochemical results would be consistent with the dramatic reduction of γ1-induced shifts of Slo1 GVs when β1 is co-expressed in HEK and LNCaP cells, they contrast with functional observations of a ternary complex with Slo1: γ1: β2 in RNA-injected Xenopus oocyte membranes. The formation of such a complex would either need to have fewer than 4 β and/or γ subunits or would require large rearrangements of their extracellular domains (relative to what we know from currently available structures, as discovered in our study and previous studies on Slo1-β4). Further electrophysiological and biochemical studies will be necessary to resolve this dichotomy.

The proposed cooperativity in assembly conferred by the LRRD domain also has implications for the previously noted all-or-none gating behavior conferred by LRRC26 on Slo1 channels^75,76^. In contrast to effects of Slo1 β subunits which incrementally shift gating as mole fraction of expressed β:Slo1 is increased^91^, increases in γ1:slo1 expressed ratio show only a changing ratio of fully shifted and unshifted population of channels^76^. Initial evaluations noted that this could arise either from two types of models: first, in which differing γ1:Slo1 stoichiometries might occur, but that a single γ1 subunit is sufficient to produce a full effect; and a second, where γ1:Slo1 assembly always involves a fixed ratio of γ1 to Slo1 subunits. However, it was also subsequently shown that single channels with an engineered construct containing only a single γ1-TM (together with its C-terminal end) segment were sufficient to produce a full gating shift (ref), but as noted above these constructs lacked an LRRD domain. The present results empirically suggest that under the conditions of expression in HEK cells the presence of the LRRC domain strongly constrains assembly to 4:4. Yet this does not preclude the earlier results supporting the idea that the presence of a single γ1-TM (together with its C-terminal end) is sufficient to produce the full gating effects.

### Plausible mechanism of LRRC26 regulation of Slo1 voltage-gating

The γ1 subunit, despite producing a larger hyperpolarizing shift in Slo1 GV than the R207A mutation, does not lead to re-orientation of the S4 gating charges. This suggests that its impact on voltage-sensor activation energetics might be lower than that of R207A. The functional effect of γ1 is thus more likely to arise from modulation of transitions that occur late in or after Slo1 voltage activation. Indeed, γ1 subunits have been proposed to profoundly enhance allosteric interactions between the voltage-sensor and pore^14^.

While the LRRD of γ1 plays a distinct and important role in the Slo1: γ1 complex assembly, the γ1-TM is critical for assembly as well as Slo1 functional regulation^14,67,72,75^. The γ1-TM kink in particular might be a critical structural element necessary for its modulatory effects. Mutation of the γ1 residue S272, localized at the kink, although it does not alter assembly, disrupts its gating shifts. The γ1-TM kink is adjacent to the S2 helix of Slo1-VSD, at the level of the hydrophobic gasket (Slo1-F160) which plays an important role in voltage-sensing. Slo1-D186 (on S3) is also positioned close to the hydrophobic gasket. These residues (F160 and D186) interact with R210 in the resting state and R213 in the activated state of the VSD. Mutation of D186 has been reported to affect voltage-sensor activation energetics but also reduce the allosteric coupling between the voltage-sensor and the PGD^37,38^. Furthermore, the γ1-TM also forms a structural contact with and slightly re-orients the pre-S1 helix. This might participate in stabilizing the bending of S4-CT, late in voltage-dependent channel opening pathway through interactions with specific residues such as F223^60^. Drawing on these previous functional results and our structure we speculate that the γ1-TM provides structural support to the non-S4 VSD helices and stabilizes them. This facilitates bending of S4-CT, late during channel activation and in effect favors channel opening (**Fig. 6**). Overall our hypothesis invokes the idea that the modulatory effects of γ1 are possibly associated with altering the energy landscape of gating, rather than inducing a specific conformational change in Slo1. Structural studies on Slo1 in complex with β4 subunits have also led to similar suggestions in the context of β4 subunit modulation of Slo1 gating^28^. More detailed functional and structural investigations will be necessary to evaluate our hypothesis. We are hopeful that the approach used here, taking advantage of mutations that uniquely impact Slo1 voltage-dependent channel opening and gating charge displacement, may unveil additional conformational intermediates that define the spatial landscape of Slo1 gating and modulation by auxiliary subunits.

**Figure 6.**
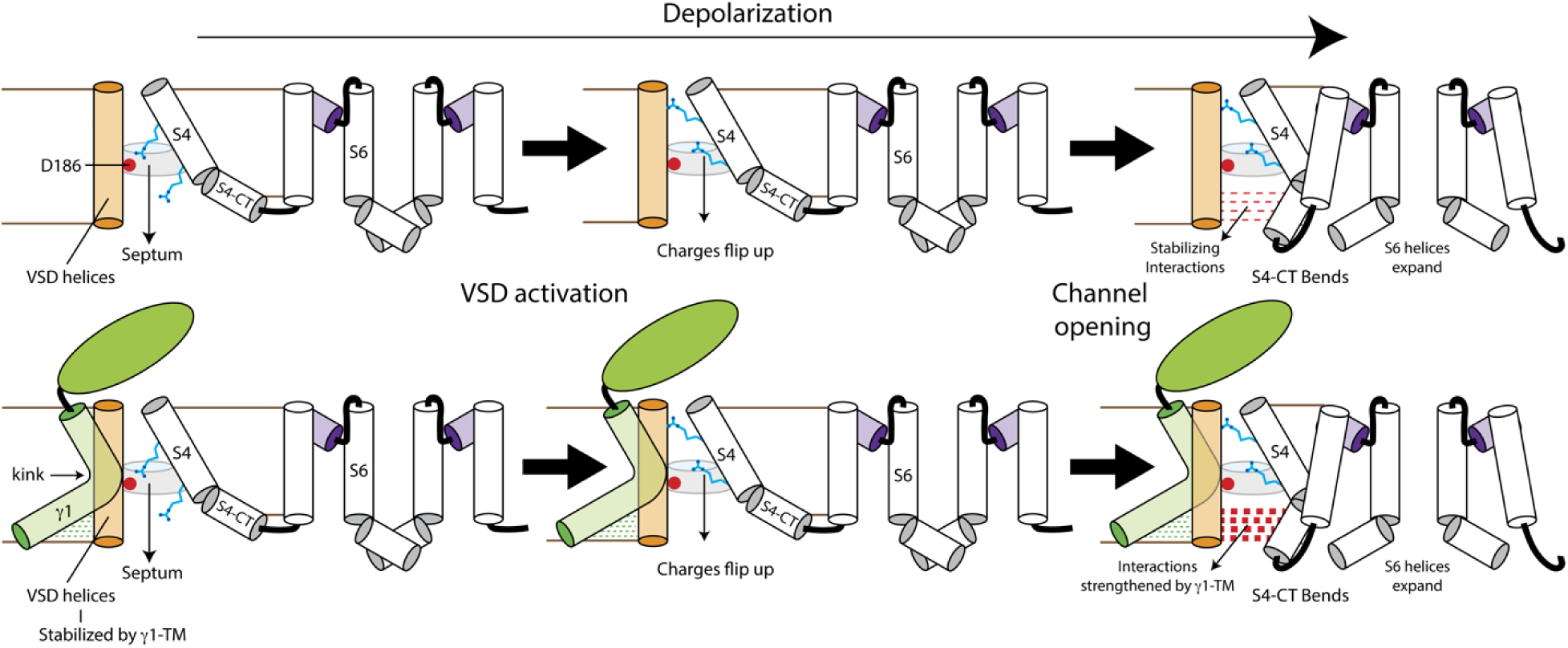
Possible mechanism underlying modulation of voltage-dependent gating by γ1 subunits. *Top*, In the absence of γ1, voltage-dependent channel opening involves a rotameric flip of the gating charges on the S4 segment which transfers the positively charged guanidium moieties of Arg residues across a focused electric field (the hydrophobic septum). Subsequently, when the S6 helices expand, as channels open, the intracellular, C-terminal end of S4 (S4-CT) rotates inward and interacts with the non-S4 helices of the VSD. These interactions (red dotted lines) might facilitate electromechanical coupling. *Bottom*, the association of γ1 subunits with Slo1 structurally stabilizes the non-S4 helices of the VSD. This strengthens the interactions between the S4-CT and non-S4 VSD helices which overall facilitates channel opening by enhancing electromechanical coupling between the VSD and the pore.

## Methods

### Expression and Purification of hSlo1 mutants

hSlo1_EM_ used in previous structural studies was generously provided by Roderick Mackinnon in the pEG vector for expression in mammalian cells. The construct featured a 57-residue deletion at the very C-terminus with respect to the native hSlo1 sequence (GI: 507922) and was followed by a 3C protease cleavage tag, eGFP and rho1D4 antibody recognition sequence. In our study we replaced this rho1D4 tag with a twin-strep tag and refer to this modified expression construct as hSlo1_EM_. All mutants were generated in the context of this construct using standard molecular biology techniques (Genscript Inc.). For protein expression, endotoxin free plasmid DNA (Qiagen) was transfected into suspension cultures of HEK293F cells, grown in Freestyle 293 media (supplemented with 2% Heat Inactivated FBS) using Linear PEI (25kDa) at a ratio of 1:3 (plasmid:PEI mass ratio). Post-transfection, cells were grown for 12-14 hrs at 37°C, and subsequently sodium butyrate was added to the transfected cells to a final concentration of 10 mM. Cultures were then transferred to 30°C and grown for another 48-54 hrs. Cells were pelleted, washed with PBS and rapidly frozen in liquid nitrogen and stored at −80°C until use. For purification, frozen cell pellets were resuspended and sonicated in ice-cold lysis buffer (500 mM KCl, 50 mM Tris, 20% glycerol, 10 mM CaCl_2_, 10 mM MgCl_2_ 1% digitonin, pH 8) and gently agitated at 4°C for 1-1.5 hrs. Total protein digitonin extracts were spun at 100,000g for 1 hr and the supernatant was incubated anti-GFP-nanobody resin (generated by PCF, University of Iowa) for 8-10 hrs. Protein bound resin was washed 4 times in batch mode, each time with 5 resin volumes of wash buffer (500 mM KCl, 50 mM Tris, 20% glycerol, 0.1% digitonin, pH 8). After the last wash, the resin was resuspended in 2x resin volume of wash buffer and incubated with Precision protease (ThermoScientific) for 12-14hrs. The eluted protein was concentrated to ∼500 μl using 100kDa centrifugation filters and the resultant protein was further purified by gel filtration chromatography. The gel filtration buffer used was 300 mM KCl, 25 mM HEPES, 0.05% digitonin, 1 mM EDTA, buffered to pH 8. 1-1.25 ml of the peak fractions of the protein was collected and concentrated to ∼6-7 mg/ml for preparing cryoEM grids. For F315A and 2D2A/4D4A mutants, ∼30 g of wet cell pellets (obtained from ∼4 L cell culture) resulted in sufficient amount of sample to prepare 6-8 EM grids. R207A expressed at a much lower level. From ∼70 g of wet cell pellets (obtained from ∼8 L cell culture) we finally obtained purified protein at ∼5 mg/ml sufficient to prepare 3-4 EM grids.

To test the effect of K^+^ ions on protein stability, hSlo1_EM_ and its mutants (D292N and Y279F) were expressed and purified similarly, except that after binding the protein to the resin, the wash buffer used (to remove contaminants and in which 3C protease cleavage was performed) was 500 mM NaCl, 2.5 mM KCl, 50 mM Tris, 20% glycerol, 0.1% digitonin, pH 8. For the FSEC runs, the running buffer was 300 mM KCl, 20 mM Tris, 0.01% L-MNG, 0.02 mM CHS, buffered to pH 8.

### Expression and Purification of hSlo1:**g**1 complex for cryoEM

The γ1 ORF (human ortholog LRRC26) used in this study was obtained from Genscript database and tagged on the C-terminus with 3C protease, mCherry and 1x FLAG tags, and finally cloned into pEG vector. hSlo1_EM_ and γ1 was co-transfected at a total plasmid weight ratio of 1:2. 1 L of HEK293F suspension culture was transfected with 2.25 mg of total DNA (0.75 mg of hSlo1_EM_ and 1.5 mg of γ1) and 6.75 ml (1 mg/ml stock) of Linear PEI (25 kDa) and expression was carried on as for hSlo1_EM_ mutants described above. For EM sample preparation, 10 L cell culture resulting in ∼95g wet cell pellet was used.

Whole cell protein extracts (extraction buffer: 500 mM KCl, 50 mM Tris, 20% glycerol, 10 mM CaCl_2_, 10 mM MgCl_2_, 1% digitonin, pH 8) were centrifuged for 1hr at 100,000g and the supernatant was incubated with streptactin resin for 12-14hrs (in batch). Protein bound resin was then washed 4 times with 5x resin volumes of Strep-wash buffer (500 mM KCl, 50 mM Tris, 20% glycerol, 0.1% digitonin, 10 mM CaCl_2_, 10 mM MgCl_2_, pH 8) and protein was eluted in Strep-wash buffer supplemented with 10 mM desthiobiotin (pH readjusted to 8 after dissolving desthiobiotin). This eluent (El1) was concentrated to ∼10 ml total volume using a 100 kDa MWCO filter and incubated with anti-Flag M2 affinity resin (Sigma) in batch for 4 hrs. The flow-through from this second affinity purification step (FT2) contained free hSlo1 tetramers. The resin bound protein was washed 4 times with 5x resin volumes of FLAG-wash buffer (500 mM KCl, 50 mM Tris, 20% glycerol, 0.1% digitonin, pH 8) and the protein was eluted in Flag-wash buffer supplemented with 3xFLAG-peptide (0.3 mg/ml). The final eluted protein (El2) was concentrated to ∼400 μl. 100 μl of commercial 3C Precision protease (ThermoScientific), supplied as ∼3 mg/ml (2 U/μl) stock in a storage buffer containing 1 mM DTT, was first volumetrically diluted 20x using FLAG-wash buffer and concentrated to <100 μl using a 30 kDa MWCO filter. This was repeated a second time until the final concentrate reached ∼3 mg/ml and 50 μl of this protease (in depleted DTT) was added to the ∼400 μl of the purified hSlo1:γ1 (eGFP, mCherry tagged) protein sample. After overnight incubation at 4°C and the hSlo1:γ1 complex (free of eGFP and mCherry tags) was further purified using gel filtration chromatography as described in the previous section. The final protein, concentrated to ∼ 6 mg/ml, was used to prepare EM grids.

### Expression and Purification of **g** subunits and mass-photometry of **g**3

The ORFs for LRRC52 (γ2), LRRC55 (γ3) and LRRC38 (γ4) were obtained from Genscript and used to generate expression constructs as described above for γ1. The proteins were expressed in transfected suspension cultures of HEK293F cells as described above. Cells expressing the individual subtypes of γ subunits were extracted in 500 mM KCl, 50 mM Tris, 20% glycerol, 1% digitonin, pH 8, clarified via ultracentrifugation and subjected to FLAG-affinity purification. After protein binding, the resin was washed 12 times with 4x resin volume of FLAG-wash buffer: 500 mM KCl, 50 mM Tris, 10% glycerol, 0.003% L-MNG, 0.02 mM CHS, pH 8. Protein was eluted in FLAG-wash buffer supplemented with 0.2 mg/ml of 3x FLAG peptide. Purified protein was first assayed using FSEC during which elution was monitored using mCherry fluorescence (Ex./Em.: 587/610 nm). The expression constructs for the LRRD domains of γ1-4 (residues 1-261 for γ1, 1-244 for γ2, 1-270 for γ3 and 1-247 for γ4) were generated similar to the full-length γ subunits and were expressed and purified identically.

For full-length γ3, concentrated purified protein (mCherry tagged) was further purified using gel filtration chromatography using FLAG-wash buffer (without glycerol). Fractions corresponding to the lower molecular weight peak were pooled, concentrated, and divided into two parts of which one was incubated with 3C protease for ∼12 hrs to release C-terminal tags. The concentrated γ3 samples (with and without the tag) were exchanged into detergent-free buffer (300 mM KCl, 20 mM Tris, pH 8) and the peak fraction was used immediately for mass-photometry experiments (Refeyn Inc). For the latter, protein was used at a final concentration of 1-3 μg/ml.

### Assembly of hSlo1 with **g**1 mutants with Dual-Color FSEC

To test the efficiency of assembly of different auxiliary subunits (and their variants, tagged as described for γ1), they were co-expressed with hSlo1_EM_ in 60 ml of HEK293F suspension cultures (plasmid weight: 45 μg hSlo1_EM_ and 90 μg auxiliary subunit and 405 μl of PEI (1 mg/ml stock)) and purified using the 2-step affinity purification described above. However, for quantitative reproducibility (as opposed to preparative biochemistry) effects of non-specific interaction with affinity chromatography resins and volumetric changes during protein purification were carefully monitored and controlled. Particularly, using 2 different variants of a model membrane protein (sea urchin ortholog of SLC9C1, or sp9C1, tagged C-terminally with eGFP-twin-strep or mCherry-FLAG), which can be purified using biochemical methods comparable to hSlo1_EM_, we determined that <5% protein was lost due to non-specific binding of tagged protein with the non-compatible resin (that is, when ∼500 μl of 1-10 μg/ml sp9C1-eGFP-twinstrep was incubated with ∼50 μl anti-FLAG resin for 4 hrs at 4°C or when ∼500 μl of 1-10 μg/ml sp9C1-mCherry-FLAG was incubated with ∼50 μl streptactin resin, >=95% protein was in the flow-through). For our assembly assay, El1 (the eluent after streptactin affinity chromatography) was concentrated to 450 μl (± 2 μl, the calibration error of our P200 pipettes). 20 μl of El1 was injected for FSEC analysis where the elution profile was simultaneously monitored by eGFP and mCherry fluorescence (Ex/Em: 488/507 nm and 587/610 nm respectively). 400 μl of El1 was incubated with 50 μl equilibrated anti-FLAG resin in spin columns. Prior to this incubation, we ensured that the resin was almost completely depleted of equilibration buffer by spinning the resin in the spin filtration columns on a benchtop centrifuge (14000 rpm for 2 mins) leaving the resin almost dry. After 4 hrs of incubation, the FLAG-resin:El2 was spinned again (at 14000 rpm for 2 mins) and the volume of FT2 was checked and ensured that it was within 390-405 μl. 20 μl of FT2 was used for FSEC analysis. The total peak height of the mCherry-based elution profile for FT2 was always <2% of that for El1, indicating almost complete capture of auxiliary subunits by the resin and thus the eGFP-based elution profile represents hSlo1_EM_ that is uncomplexed with the co-expressed auxiliary subunits. The eluent from the second affinity step (El2) was also analyzed by FSEC and the eGFP and mCherry fluorescence intensities of the sample peak should be proportional to the stoichiometry of the purified complexes. However, weak FRET between the fluorescent tags could potentially obfuscate stoichiometric inferences. Hence, the purified protein was treated with 3C protease to release the fluorescent tags and the resultant samples were also retested by FSEC. The eGFP:mcherry intensity ratio (integral of the peaks) of cleaved fluorescent tags, ρ_GC_, (which have a retention volume 4-5 ml lower than when they are attached to the complexes) retain the stoichiometric information of the purified protein but is FRET-free. To calibrate ρ_GC_ to a defined stoichiometry, we used sp9C1 which is an obligate homodimer. sp9C1-eGFP-twinstrep and sp9C1-mCherry-FLAG was co-expressed in HEK293F cells and purified using the 2-step affinity scheme. The final purified protein, El2, necessarily contains 1 copy of eGFP and 1 copy of mCherry and its ρ_GC_ defined the calibrated ρ_GC_ for 1:1 complexes. Furthermore, commercially obtained purified free-eGFP and free-mCherry (Abcam) were mixed at different molar ratios, analysed using FSEC, and used to calibrate ρ_GC_ for various other stoichiometries. Total expression of all γ variants when co-expressed with hSlo1_EM_, as quantified by measurement of total mCherry fluorescence in the whole cell detergent extracts, were all within ± 10% of each other.

To quantify fractional assembly we performed Gaussian analysis of the FSEC profiles. The eGFP profiles of various fractions were fitted to Gaussian curves and the corresponding mCherry profiles were used to validate and check the accuracy of the Gaussian parameters. For the analysis, the difference chomatogram (Diff) was generated by subtracting FT2 from El1. This corresponds to Slo1-auxiliary subunit complex. Diff, after peak normalization, was overlayed on Peak Normalized El2 to ensure that the profiles (peak positions and width) matched well. Unnormalized Diff was fitted to a sum of 2 Gaussians: 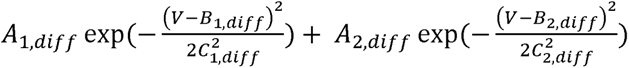, where the first component corresponds to the main/desired complex of tetrameric hSlo1 and the auxiliary subunit and the second component corresponds to other higher molecular weight aggregates (if present they are left shifted with respect to the first by ∼1.5 ml on a Superose 6 Increase 10/300 column). FT2 (eGFP profile) was fitted to a sum of 3 Gaussians: 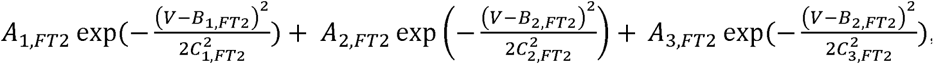, where the first gaussian corresponds to free Slo1 tetramers, the second accounts for some molecular aggregates of Slo1 (which may be present but is always <10% of total Slo1), and the third corresponds to lower molecular weight peaks of disassembled Slo1 (which usually is 20-40% of total Slo1). Fractional assembly (amount of tetrameric hSlo1_EM_ in complex with auxiliary subunit relative to total tetrameric Slo1_EM_) was quantified as: 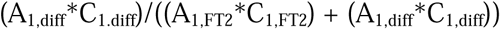

### CryoEM grid preparation and imaging

The purified and concentrated proteins were incubated with an additional 5 mM EDTA for 2-4 hrs before preparation of EM grids. 3.5 μl of concentrated purified protein was applied to glow-discharged copper holey carbon grids (Quantifoil R 1.2/1.3, 300 mesh). In all cases, grids were blotted at 4°C for 5 s at 100% humidity with a blot force of 0 and then plunge frozen in liquid ethane using a Vitrobot Mark IV (Thermofisher Scientific). All data were collected on a Titan Krios (operating at an accelerating voltage of 300kV), equipped with a K3 Detector (Gatan). Images were recorded with EPU software (ThermoFisher Scientific) in super-resolution mode with a pixel size of 0.54 Å, and a nominal defocus of −0.9 to −1.7µm. For all datasets, a total dose of 72 electrons/Å^2^ fractionated over 40 frames was used.

### Image Processing and Map Calculation

Image processing and map calculations were performed using CryoSPARC^92^ and Relion^93^. Motion corrected (via Patch motion correction) were subjected to contrast transfer function (CTF) estimation (Patch CTF). Micrographs with CTF resolution > 6Å or total pixel drift > 60 pixels were discarded. Blob picker was used to pick particles from 500-1000 images, which were subject to several rounds of reference-free 2D class averaging. The final set of 2D projections (3-10) were used for template-based particle picking from curated movies. Particles corresponding to these classes (20-50K) particles were used to obtain a representative initial 3D map of the protein via Ab initio reconstruction (1 class). The template picked particles were subject to several rounds of heterogenous refinements using the initial 3D map as reference. At each step of heterogenous refinement, the particles which were classified to low resolution classes were discarded. The set of particles finally retained were refined using Non-uniform, CTF (local and global) and local (with mask around the full protein) refinement routines on CryoSPARC (with C4 symmetry imposed) to obtain the final maps (for F315A and 4D4A/2D2A datasets). In the R207A dataset, the particles were exported to Relion and further classified using 3D classification, without alignment. The optimal set of particles were imported back into CryoSPARC and subject to Non-uniform and CTF refinements to obtain the final map. For the hSlo1:γ1 complex data processing initially resulted in a map where all relevant densities were clearly observed but the extracellular LRRD density for γ1 was relatively lower in quality (hSlo1-γ1-Full map). In separate workflows, these particle stacks were subject to particle subtraction (on Cryosparc) to remove density for the LRRD and for the gating-ring and subsequent refinements (using masks around the non-LRRD or the non-gating-ring parts of the complex respectively) with C4 symmetry imposed were used to obtain the hSlo1-γ1-LRRD-masked and hSlo1-γ1-GR-masked maps. For the F315A dataset, particle stacks were binned by a factor of 1.667 while for the others they were binned by a factor of 2.

### Model Building and Refinement

The DVF model of hSlo1_EM_ (6V3G) was used to build atomic models for the hSlo1_EM_ mutants into our density maps. For γ1, an alphafold predicted model was used to guide model building. For F315A, the sharpened map was used for model building but the rest of the structural models were built into the unsharpened maps. Models were built via iterative rounds of manual model building on COOT and real space refinement in Phenix. The final refined atomic models were validated using MolProbity^94^. All structural analyses were performed on UCSF Chimera. Structural figures were generated using UCSF ChimeraX or Pymol.

### Electrophysiology and Data Analysis

As described before^95^, Slo1 channels were expressed in stage IV Xenopus oocyte by cRNA injection of mouse Slo1 (mSlo1) or human Slo1 (hSlo1) without or with human LRRC26 (hγ1). Macroscopic and single channel currents were recorded in the inside-out patch configuration with an Axopatch 200B amplifier (Molecular Devices, Sunnyvale, CA). Data were acquired with the Clampex program from the pClamp software package (Molecular Devices, San Jose, CA). The pipette resistance for ionic current recording was typically 1-2 MΩ (macroscopic) or 4-5 MΩ (single) after heat polishing. The pipette solution which bathes the extracellular face of patch membranes) contained (in mM): 140 K-methanesulfonate (KMES), 20 KOH, 2 MgCl2, 10 HEPES. Solutions applied to the cytosolic side of the membrane contained (in mM): 140 KMES, 20 KOH, 10 HEPES. 5 mM HEDTA was used for 10 μM Ca^2+^ and 5 mM EGTA for 0 μM Ca^2+^ cytosolic solutions. The pH of all solutions was adjusted to 7 with methanesulfonic acid. An SF-77B fast perfusion stepper system (Warner Instruments, Hamden, CT) was used to produce solution exchange at the tip of the recording pipette.. Experiments were performed at room temperature (∼22-25 °C). All chemicals were purchased from Sigma-Aldrich (St. Louis, MO).

Tail current amplitude measured 150 μs after repolarization to −120 mV was used to define the G-V relationship of Slo1 channels. G-V curves were fit by the Boltzmann function: (*G*= (*G_max_* /(1 + *exp(-z(v - v_h_)/kr)*). G_max_ is maximal conductance, z is apparent voltage-dependence in units of elementary charge, V_h_ is the voltage of half-maximal activation, and k and T have their usual physical meanings. The open probability of Slo1 single channel current was determined by the threshold-based Event Detection method in Clampfit 9.2 (Molecular Device). Data were analyzed using OriginPro 7.5 (OriginLab Corporation) or Clampfit 9.2. Error bars in the figures represent SDs.

### DATA AVAILABILITY

All experimental data will be made available upon reasonable request. Sharpened, unsharpened and half-maps for 6 reconstructions and 5 atomic models (described in Supplementary Tables 1 and 2) have been deposited to EMDB/PDB repositories.

## ACKNOWLEDGEMENTS

This research was supported by grants to S.C. from NIH (R01-GM145719) and Department of Molecular Physiology and Biophysics, University of Iowa and NIH grant (R35-GM118114) to C.L.. We thank Dr. Vera Moiseenkova-Bell and acknowledge the use of instruments at the Beckman Center for Cryo-Electron Microscopy at the University of Pennsylvania Perelman School of Medicine. We thank Dr. Stefan Steimle for assistance with Krios microscope operation at the Beckman Center for Cryo-Electron Microscopy at the University of Pennsylvania, Perelman School of Medicine. We thank Dr. Vivian Gonzalez-Perez (Washington University at St. Louis) and Dr. Alexandria N. Miller (University of Iowa) for helpful discussions, Dr. Zhen Xu (PCF, University of Iowa) for providing anti-GFP conjugated resin and Sankar Baruah (PCF, University of Iowa) for help with gel filtration chromatography and mass-photometry

## AUTHOR CONTRIBUTIONS

S.C. designed and directed research. G.S.K performed biochemistry experiments. K.P. built atomic models. S.C. assisted with biochemistry experiments, analyzed biochemical data and performed single-particle analysis. Y.Z. performed electrophysiology experiments. Y.Z. and C.L. analyzed electrophysiology data. S.C. wrote initial draft of manuscript and modified it together with C.L. All authors contributed towards preparing manuscript figures.

