## Supplementary Figure for "High-resolution structures illuminate key principles underlying voltage and LRRC26 regulation of Slo1 channels"

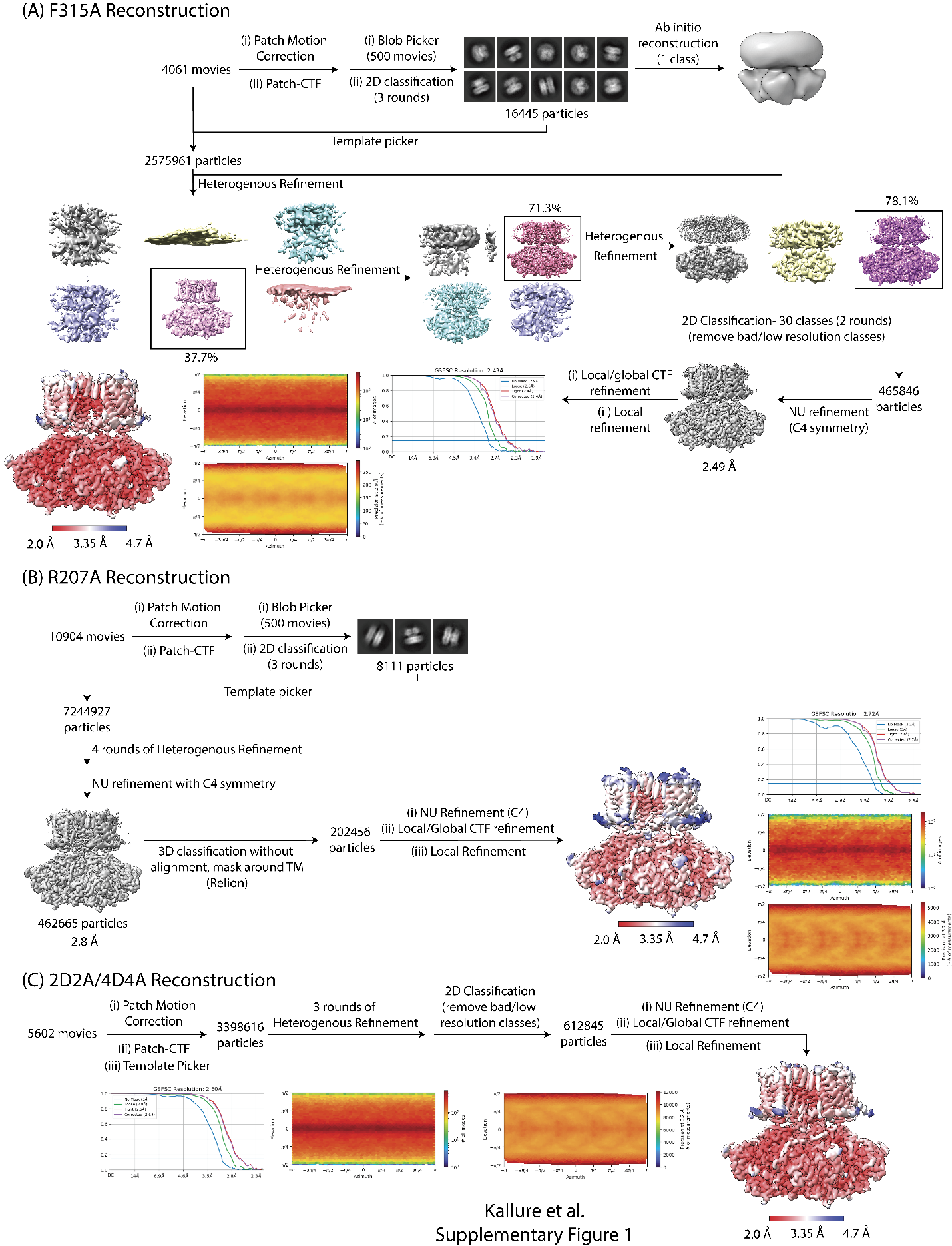

**Supplementary Figure 1. Single Particle Reconstruct of hSlo1 mutants in 5mM EDTA.** Overview of reconstruction workflow for hSlo1 mutants F315A **(A)**, R207A **(B)** and 2D2A/4D4A **(C)**. In each case the FSC-curves and the orientation distribution plots are provided for the unsharpened, final map (colored according to local resolution estimation FSC = 0.5)

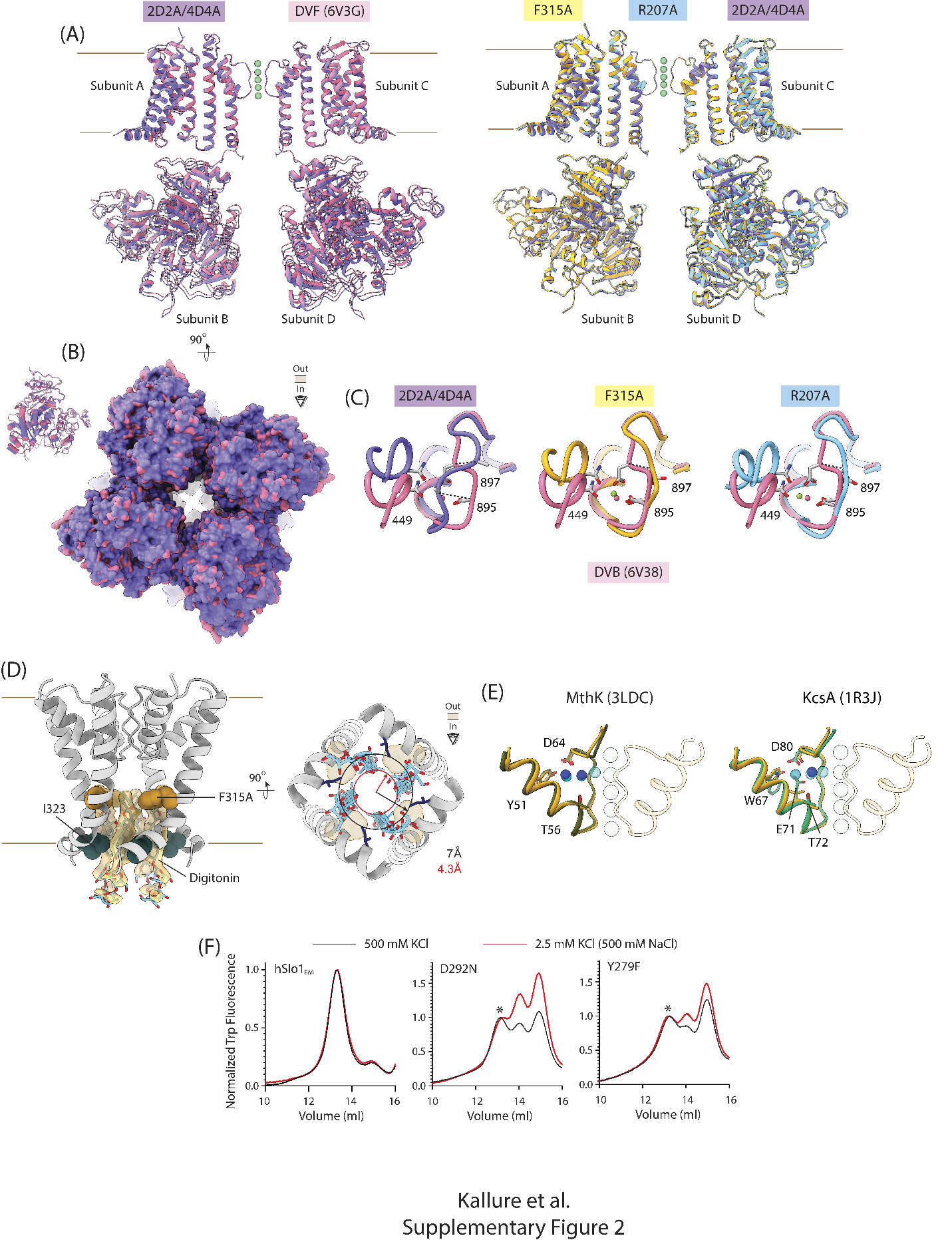

**Supplementary Figure 2. Structural features of hSlo1 mutants. (A)** *Left,* Overlay of the structural models of 2D2A/4D4A (purple) with the previous Divalent Free state model of hSlo1 (6V3G; magenta), aligned at the selectivity filter. Green spheres represent the K^+^ ions. The transmembrane domains of 2 diagonally apposed subunits are shown. The cytoplasmic domains shown belong to the 2 other subunits. *Right,* Structural overlay of F315A (yellow), R207A and 2D2A/4D4A aligned at the selectivity filter shown in a similar view as in the left panel. **(B)** Bottom-up view of the cytoplasmic gating rings of F315A and 6V3G aligned and colored as in (A), in surface representation. Inset shows a single cytoplasmic domain with the helices of RCK1 in ribbons and those of RCK2 domain in tubes. **(C)** Close up view of the Calcium bowl locus of 2D2A/4D4A, F315A and R207A (colored as in (A)) overlayed with the previous Divalent Bound state (6V38) in magenta, showing the critical sidechains coordinating the ion density. The sidechain orientation and Cα position of residue D897 is markedly different in our structures relative to the DVB state. In each of the 3overlays, magenta spheres represent the Ca^2+^ density in the DVB state, while the green spheres show the ion-like density observed in F315A and R207A maps (possibly representing a K^+^ ion). In 2D2A/4D4A no such ion density was observed. **(D)** *Left*, the selectivity filter and the S6 helices (in side view) with the occluding detergent molecules (blue/red sticks) shown in context of the final model for 2D2A/4D4A. Densities for the occluding detergent molecules are shown in yellow. Residues F315 and I323 are shown as spheres. *Right,* bottom-up of the pore showing the reduction of the access pathway at the inner mouth of the pore (at the level of I323 (dark blue sticks)), caused by the detergent molecules. Transparent dark yellow spheres represent F315 residue. Detergent binding sites to membrane proteins could correlate with lipid (or small hydrophobic molecule) interaction sites, raising the possibility that under specific conditions, even in native membranes the inner pore of hSlo1 can become occupied by specific lipids which would likely alter its gating or ion conduction. The densities for these digitonin molecules are dramatically attenuated in the F315A map which could indicate that this hydrophobic residue influences the interaction of Slo1 with hydrophobic small molecules in the inner pore. Of note, a recent reconstruction of the MthK channel^43^ (ref), an ancient and widely studied Ca^2+^ activated channel related to Slo1, has shown that the residue corresponding to F315 (F87 in MthK) interacts with the hydrophobic groups of the pore blocker, bbTBA. **(E)** Comparison of the selectivity filter and P helices of 2 adjacent subunits in F315A model (yellow) and MthK (PDB:3LDC, left) and KcsA (PDB-IDL:1R3J, right). In each case, dark blue spheres represent the water molecules in the structures of the prokaryotic K^+^ channels. Light blue/transparent spheres are the water molecules in F315A. Critical residues involved in direct H-bonding or water-bridged H-bond networks are shown in sticks and numbered according to MthK/KcsA structures. In MthK the residues are identical to hSlo1 but in KcsA, W67 and E71 are different (correspond to Y279 and V284 in hSlo1). **(F)** Size-exclusion chromatography profiles of hSlo1 and its mutants, purified in 500 mM KCl buffers or 500 mM NaCl/2.5 mM KCl buffers, monitored via intrinsic tryptophan fluorescence of purified proteins. The * mark indicates the tetrameric species in D292N (middle) and Y279F (right).

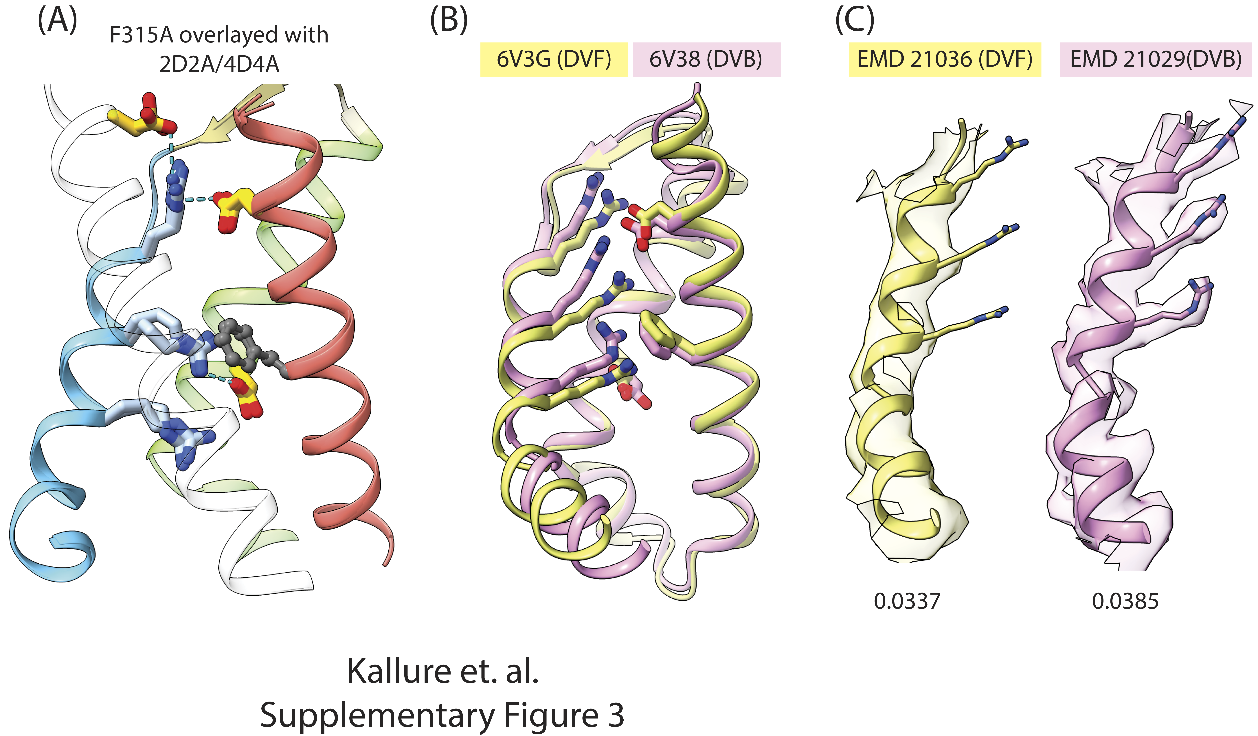

**Supplementary Figure 3. Voltage-sensing charges in different Slo1 structures. (A)** Structural overlay of the voltage-sensors of F315A and 2D2A/4D4A show an almost exact overlap at the backbone level. The arginines R207, R210 and R213 (in blue sticks) on S4 (blue ribbon) show a similar orientation, relative to the acidic residues on S1 (white transparent ribbon), S2 (salmon ribbon), S3 (green ribbon) and F160 on S2. **(B)** Structural overlay of voltage-sensors of the previous DVF (6V3G), yellow, and DVB (6V38), pink, states. (S1 helix is not shown). **(C)** Density maps and the orientation of arginine sidechains of S4 in DVF (yellow) and DVB (pink) states contoured at similar thresholds (shown below)

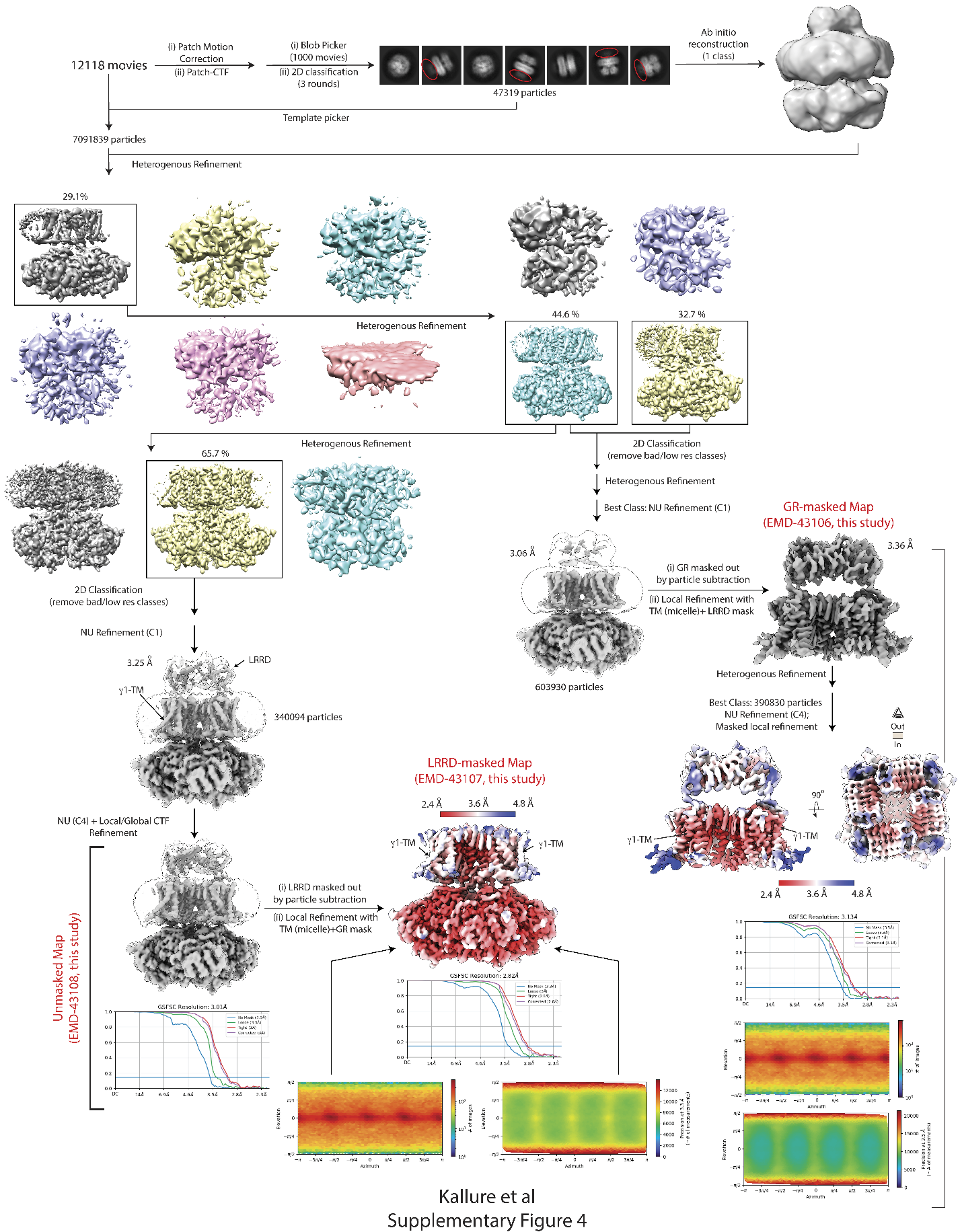

**Supplementary Figure 4. Reconstruction workflow of hSlo1:hγ1 complex in presence of EDTA.** FSC maps and orientation distribution plots for the unmasked, LRRD-masked, and GR-masked maps are specifically marked. The final unmasked map is shown at 2 different contour levels – low threshold view (in white) shows the outline of the micelle and the LRRD ring while the high threshold view (in gray) shows the densities of TM and RCK domains clearly, but the LRRD density appears fragmented. Unsharpened LRRD-masked and GR-masked maps are colored according to local resolution estimates (FSC = 0.5)

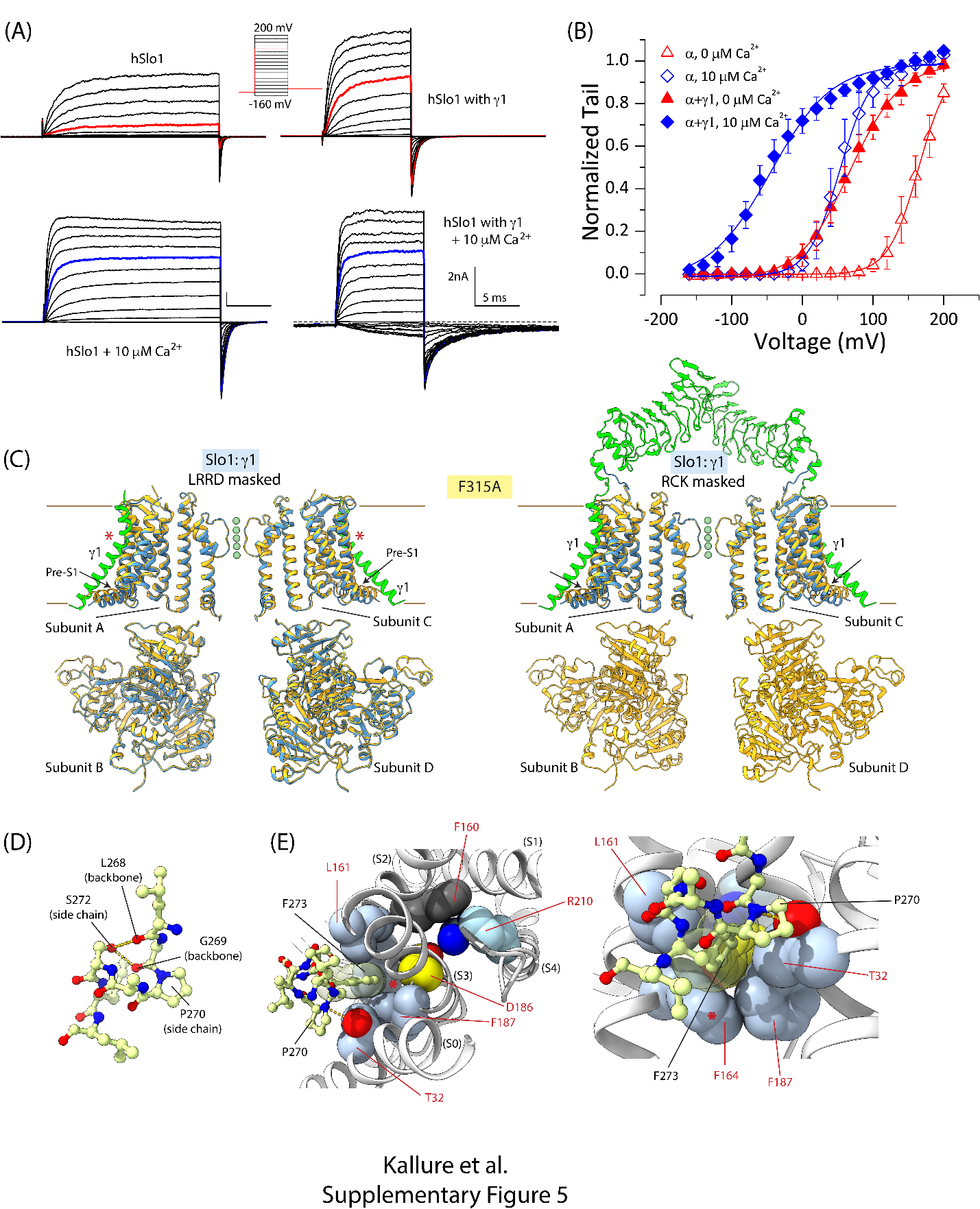

**Supplementary Figure 5. Function and structure of hSlo1:γ1. (A)** Macroscopic currents of hSlo1 (left column) or hSlo1+hγ1 (right column) evoked by a voltage protocol shown in the inset in either 0 (top row) or 10 (bottom row) μM [Ca^2+^]_in_. Channels were expressed in Xenopus oocytes. **(B)** The G-V relationships of hSlo1 and hSlo1+hγ1 determined from tail currents. The solid lines are Boltzmann fit: hSlo1, 0 μM [Ca^2+^]_in_: 164 mV, 1.17*e*; hSlo1, 10 μM [Ca^2+^]_in_: 53 mV, 1.13*e*; hSlo1+hγ1, 0 μM [Ca^2+^]_in_: 67 mV, 0. 70e; hSlo1+hγ1, 10 μM [Ca^2+^]_in_: -45 mV, 0.60e. Note that the GV curves with γ1 can be somewhat better fit with a double Boltzmann reflecting that some fraction of BK channels lack γ1 subunits. **(C)** Overlay of the structural models of F315A (yellow) with Slo1-γ1 complex (Slo1 in blue and γ1 in green) built into the LRRD-masked map (*left*) and the GR-masked map (*right*). The transmembrane domains of diagonally apposed subunits together with their partnering γ1 subunits are shown. The cytoplasmic domains belong to the 2 other diagonally arranged subunits. Overall, Slo1 models superposed very well except at the pre-S1 helix (marked with arrows) which is displaced downward in the Slo1:γ1 models. The kink in γ1 is marked by a red asterisk. **(D)** Close up view of the backbone and side-chains of the residues in the γ1-TM kink locus. The sidechain of S272 is within 3 Å of backbone carbonyls of L268 and G269, probably engaged in H-bonds. **(E)** Juxtaposition of the γ1-TM kink (ball and stick representation) with Slo1 voltage-sensor in a top-down view (*left*) and a tilted side-view (*right*). F273 fits into a hydrophobic pocket formed by L161, F164, F187 and T32, just “behind” the critical D186 and F160 residues. The latter face into the VSD core and engage the gating charge, R210, in the resting state. The red asterisk marks the Slo1 residue F164. (Slo1 and γ1 residues are marked in red and black, respectively. Slo1 VSD helices are marked in parentheses).

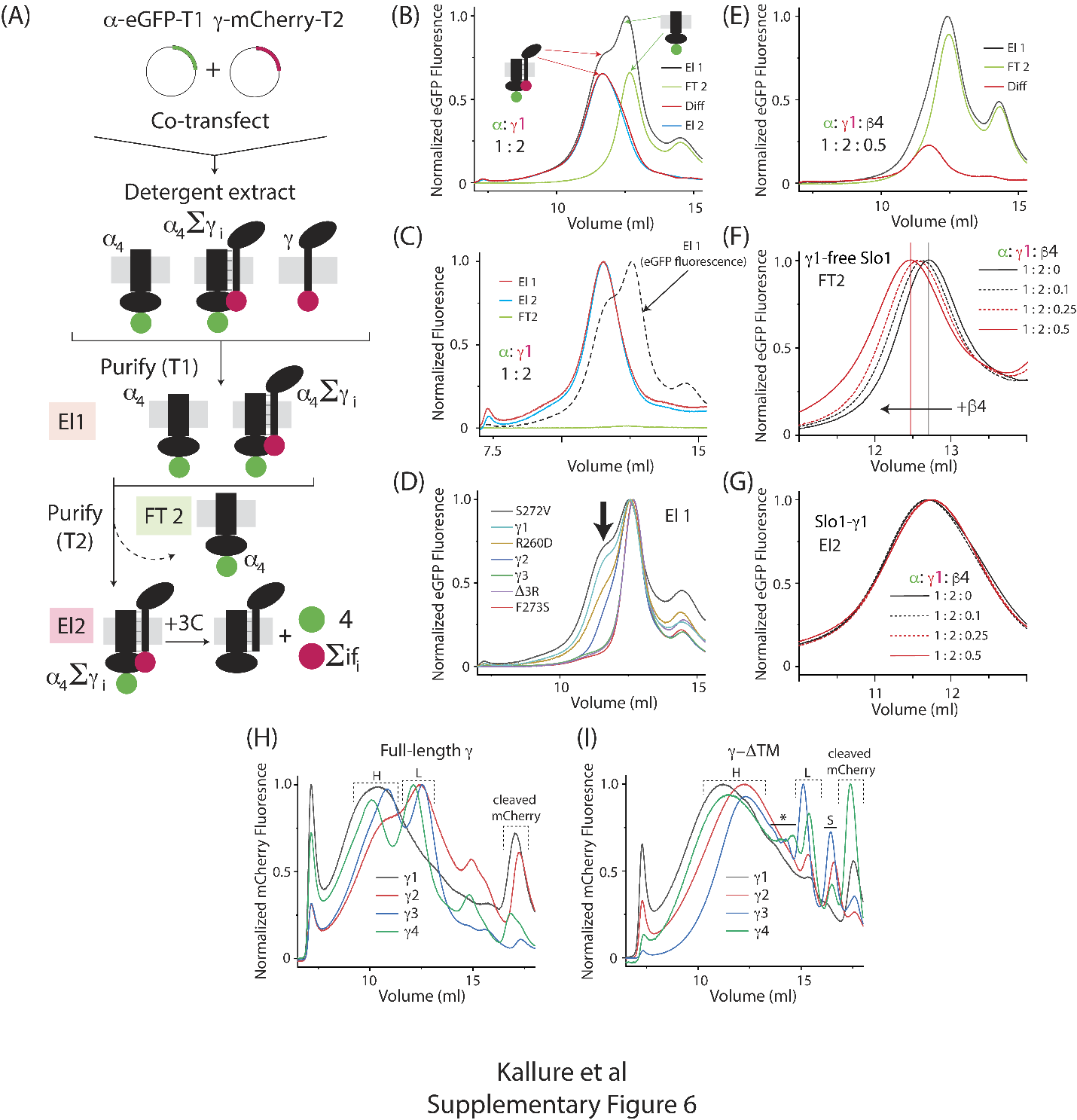

**Supplementary Figure 6. Assembly of Slo1:γ1. (A)** The 2 step purification scheme used to test assembly of Slo1 and auxiliary subunit variants as described in Methods. hSlo1 is conjugated to eGFP and purification tag, T1 (twin-strep tag), while the auxiliary subunits are tagged with mCherry and FLAG (T2). $\alpha_{4}\sum\gamma_{i}$ implies complexes of tetrameric hSlo1 and varying (i=1, 2, 3 or 4) copies of the γ1 subunits that may exist in cells/cellular extracts. After 2 step affinity purification, treatment with protease (3C) releases the fluorescent tags (eGFP and mCherry) in the ratio of the number of copies of hSlo1 subunit (4) in the complex to the average number of copies of γ1 subunit in the complex ($\sum_{i=1}^{4} if_{i}$, where f_i_ is the fraction of the complex with ‘i’ copies of γ1). **(B)** Representative size exclusion chromatography profiles of El1, El2 and FT2 as described in (A) for hSlo1:WT-hγ1, monitored by eGFP fluorescence. El1 and FT2 are normalized to the highest peak of El1 which corresponds to γ-free tetrameric Slo1. Diff represents the difference of El1 and FT2 (numerical subtraction) representing the profile of γ-complexed Slo1. El2 (which corresponds to the experimental profile of the doubly purified hSlo1:hγ1 complex) is normalized to the peak of Diff and overlaps well with Diff profile. **(C)** Representative size exclusion chromatography profiles for El1, El2 and FT2 for hSlo1:WT-hγ1 complex monitored by mCherry fluorescence, all normalized to the peak height of El1 showing the complete capture of all γ-containing complexes at the second affinity purification step (FT2) and the overlap of the γ-containing complexes between the first and second affinity purification steps (El1 and El2). The El1-eGFP profile (as in (B)) is also shown as a dotted line. **(D)** Size exclusion chromatography profiles of El1 (total hSlo1 containing protein isolated) obtained upon co-expression of hSlo1 with different mutants of hγ1. All profiles are dominant peak normalized. Arrow indicates the species corresponding to hSlo1:γ1 complex in each case. **(E)** Representative size exclusion chromatography profiles of El1 and FT2 obtained upon co-expression of hSlo1, hγ1 and hβ4 (at the indicated transfection ratio). The relative amount of γ-containing complexes, as indicated by Diff, decreases significantly relative to (B). **(F), (G)** At various levels of β4 expression (different amounts of transfected DNA), the size exclusion chromatography profiles for γ1-free complexes (FT2) shift left (indicating increase in species size/hydrodynamic radius) with increasing amount of β4 expression/transfection, but the profiles for the γ1-containing complexes (El2) **(G)** are almost superimposable. **(H)** Size exclusion chromatography profiles for affinity purified full-length **(H)** and LRRD domains (Δ-TM) **(I)** of mCherry-tagged γ subunits, monitored by mCherry fluorescence. In both cases, a large/broad oligomeric peak (marked as H) and a lower oligomeric peak (L) were observed. For full-length γ3, the species analyzed by mass photometry corresponds to ‘L’. Multiple, heterogenous oligomeric species are observed in the Δ-TM variants (marked as * or ‘S’).

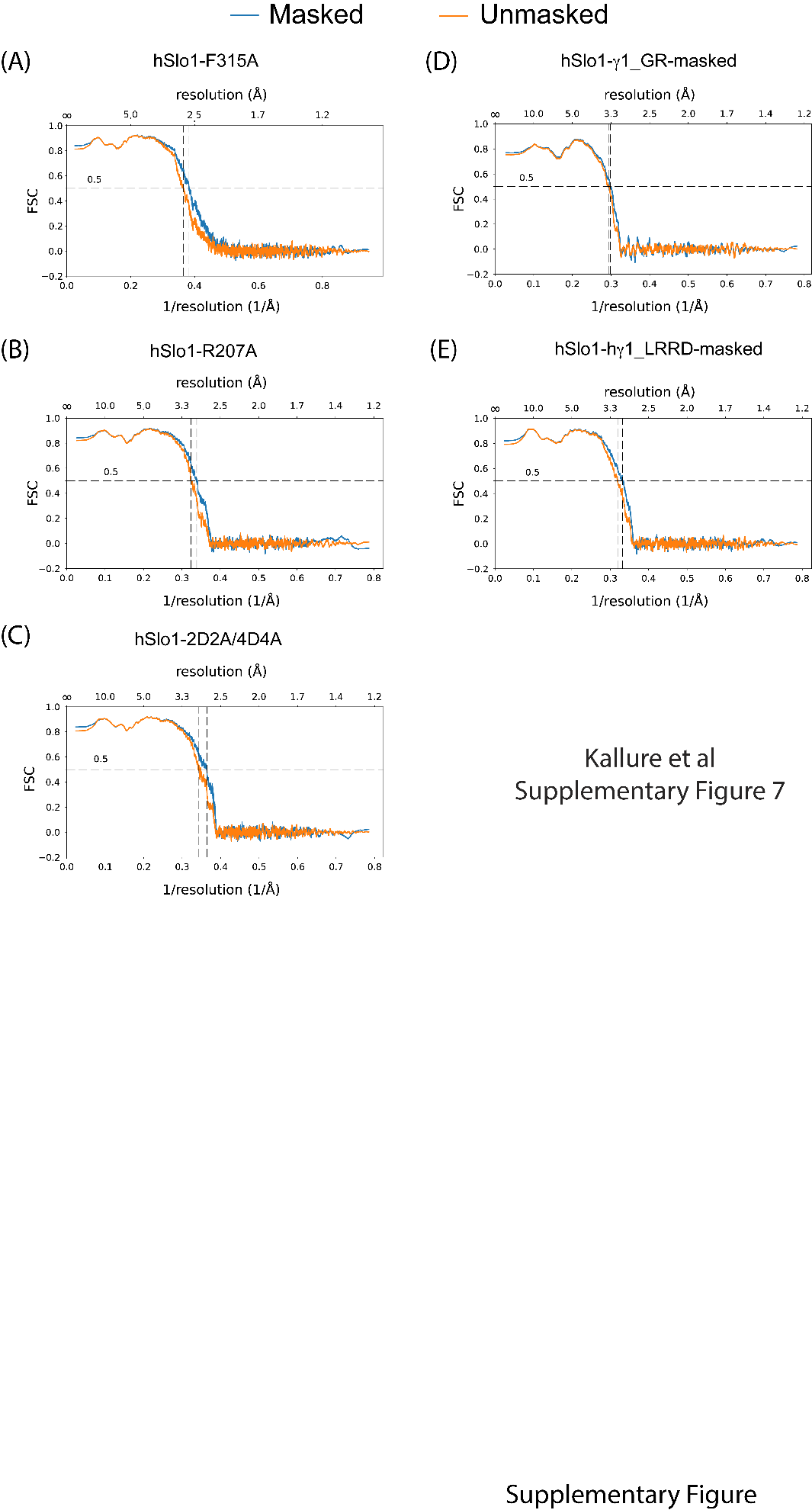

**Supplementary Figure 7. Map-to-model FSC curves.** Map-to-model FSC validation curves for the 5 models that have been generated in this study. No model was built into the unmasked hSlo1:γ1 map.

**Table I. Cryo-EM data collection, refinement and validation statistics for hSlo1-mutant reconstructions**

|  | **F315A (C)** | **R207A (C)** | **2D2A/4D4A (C)** |
| --- | --- | --- | --- |
| EMDB # | 42986 | 42988 | 42989 |
| PDB-ID | 8V60 | 8V63 | 8V64 |
| **Data collection and processing** |  |  |  |
| Magnification | 81000 | 81000 | 81000 |
| Voltage (kV) | 300 | 300 | 300 |
| Electron exposure (e–/Å^2^) | 72 | 72 | 72 |
| Defocus range (μm) | -0.9 to -1.7 | -0.9 to -1.7 | -0.9 to -1.7 |
| Pixel size (Å) (super-resolution)  (Binned by 2 during data processing: except F315A) | 0.54  (Binned by 1.667) | 0.54 | 0.54 |
| Symmetry imposed | C4 | C4 | C4 |
| Initial particle images | 2575961 | 7244927 | 3398616 |
| Final particle images (no.) | 465846 | 202456 | 612845 |
| Map resolution (Å) | 2.43 | 2.72 | 2.60 |
| FSC threshold | 0.143 | 0.143 | 0.143 |
| **Refinement** |  |  |  |
| Model resolution (Å) | 2.6 | 3.0 | 2.8 |
| FSC threshold | 0.5 | 0.5 | 0.5 |
| Map sharpening *B* factor (Å^2^)* | -50 (-83.6) | (-86.4) | (-98.2) |
| **Model Composition** |  |  |  |
| Non-Hydrogen atoms | 31464 | 31665 | 31541 |
| Protein Residues | 3624 | 3624 | 3640 |
| Ligand | 77 | 81 | 73 |
| ***B-factors (Å^2^)*** |  |  |  |
| Protein | 73.16 | 145.00 | 125.00 |
| Ligand | 140.29 | 202.67 | 178.67 |
| ***R.m.s Deviations*** |  |  |  |
| Bond lengths (Å) | 0.004 | 0.003 | 0.002 |
| Bond Angles (°) | 0.590 | 0.574 | 0.548 |
| ***Validation*** |  |  |  |
| MolProbity Score | 1.15 | 1.19 | 1.08 |
| Clash score | 3.63 | 4.04 | 2.89 |
| Poor rotamer (%) | 0.00 | 0.00 | 0.00 |
| ***RamaChandran*** |  |  |  |
| Favored (%) | 98.27 | 98.33 | 98.36 |
| Allowed (%) | 1.73 | 1.67 | 1.64 |
| Outlier (%) | 0.00 | 0.00 | 0.00 |

*B-factors auto-calculated by final refinement routines in parentheses. For F315A, B-factor of -50 was used to sharpen the final map.

**Table II. Cryo-EM data collection, refinement and validation statistics for hSlo1-G1 reconstructions**

|  | **Slo1-γ1 (GR-Masked)** | **Slo1-γ1 (LRRD Masked)** | **Slo1-γ1 Unmasked** |
| --- | --- | --- | --- |
| EMDB # | 43106 | 43107 | 43108 |
| PDB-ID | 8VAV | 8VAZ |  |
| **Data collection and processing** |  |  |  |
| Magnification | 81000 | 81000 | 81000 |
| Voltage (kV) | 300 | 300 | 300 |
| Electron exposure (e–/Å^2^) | 72 | 72 | 72 |
| Defocus range (μm) | -0.9 to -1.7 | -0.9 to -1.7 | -0.9 to -1.7 |
| Pixel size (Å) (super-resolution) (Binned by 2 during data processing) | 0.54 | 0.54 | 0.54 |
| Symmetry imposed | C4 | C4 | C4 |
| Initial particle images | 7091839 | 7091839 | 7091839 |
| Final particle images (no.) | 390830 | 340094 | 340094 |
| Map resolution (Å) | 3.13 | 2.82 | 3.01 |
| FSC threshold | 0.143 | 0.143 | 0.143 |
| **Refinement** |  |  |  |
| Model resolution (Å) | 3.4 | 3.0 |  |
| FSC threshold | 0.5 | 0.5 |  |
| Map sharpening *B* factor (Å^2^) | (-119.6) | (-104.3) | (-112.7) |
| Model Composition |  |  |  |
| Non-Hydrogen atoms | 18532 | 29404 |  |
| Protein Residues | 2096 | 3704 |  |
| Ligand | 64 | 4 |  |
| ***B-factors (Å^2^)*** |  |  |  |
| Protein | 152.14 | 128.48 |  |
| Ligand | 160.03 | 99.77 |  |
| ***R.m.s Deviations*** |  |  |  |
| Bond lengths (Å) | 0.002 | 0.002 |  |
| Bond Angles (°) | 0.668 | 0.502 |  |
| ***Validation*** |  |  |  |
| MolProbity Score | 1.21 | 1.19 |  |
| Clash score | 4.28 | 4.00 |  |
| Poor rotamer (%) | 0.00 | 0.00 |  |
| ***RamaChandran*** |  |  |  |
| Favored (%) | 98.31 | 98.50 |  |
| Allowed (%) | 1.69 | 1.50 |  |
| Outlier (%) | 0.00 | 0.00 |  |

No model was built or deposit for the Slo1-γ1 unmasked map.
